# A top-down slow breathing circuit that alleviates negative affect

**DOI:** 10.1101/2023.02.25.529925

**Authors:** Jinho Jhang, Shijia Liu, David D. O’Keefe, Sung Han

## Abstract

Breathing is profoundly influenced by both behavior and emotion^1–4^ and is the only physiological parameter that can be volitionally controlled^4–6^. This indicates the presence of cortical-to-brainstem pathways that directly control brainstem breathing centers, but the neural circuit mechanisms of top-down breathing control remain poorly understood. Here, we identify neurons in the dorsal anterior cingulate cortex (dACC) that project to the pontine reticular nucleus caudalis (PnC) and function to slow breathing rates. Optogenetic activation of this corticopontine pathway (dACC→PnC neurons) in mice slows breathing and alleviates behaviors associated with negative emotions without altering valence. Calcium responses of dACC→PnC neurons are tightly correlated with changes in breathing patterns entrained by behaviors, such as drinking. Activity is also elevated when mice find relief from an anxiety-provoking environment and slow their breathing pattern. Further, GABAergic inhibitory neurons within the PnC that receive direct input from dACC neurons decrease breathing rate by projecting to pontomedullary breathing centers. They also send collateral projections to anxiety-related structures in the forebrain, thus comprising a neural network that modulates breathing and negative affect in parallel. These analyses greatly expand our understanding of top-down breathing control and reveal circuit-based mechanisms by which slow breathing and anxiety relief are regulated together.

## Main Text

Breathing is more than simply a respiratory response to maintain gas homeostasis. Breathing rhythms are entrained by various purposeful (swallowing, vocalization) and emotional (sighing, yawning, purring) behaviors that require orofacial and pharyngeal activity^1,5,7,8^. Moreover, humans can control breathing rhythms via conscious effort – volitional breathing^4,6,9^ – and slow breathing or mindfulness skills are practiced across cultures to regulate emotion^1,5,10,11^. Given this knowledge, the primary motor cortex^12^ and high-order cortical regions, including the anterior cingulate cortex (ACC)^6,13,14^ and the supplementary motor area^12,15^, have been implicated in the central, non-respiratory control of breathing. However, circuit-level research on breathing control has primarily focused on the medullary respiratory groups and their roles in generating rhythmic patterns of breathing. Precise mechanisms for the cortical control of breathing, in particular how top-down breathing circuits affect brainstem networks, remain elusive^4,5^. In this study, we identify a prefrontal-to-pontomedullary breathing circuit that slows breathing and alleviates negative affect in parallel.

### Identification of a top-down neuronal circuit that slows breathing

To identify circuits that could potentially relay top-down cortical inputs to breathing centers in the brainstem, we performed an extensive search of the Mouse Brain Connectivity Atlas (using a ‘target search’ tool; Allen Brain Institute, connectivity.brain-map.org) and the Mouse Brain Atlas (mouse.brain-map.org) in the Allen institute for Brain Sciences (Extended data Fig. 1). We looked for regions that: 1) receive inputs from the prefrontal cortex, 2) project to pontomedullary breathing centers, and 3) are enriched for markers of inhibitory cell types (for inhibitory regulation of breathing). One of several regions that satisfies these three criteria (Extended data Fig. 1a-d) was the pontine reticular nucleus caudalis (PnC). To identify prefrontal neurons that project to the PnC, we performed retrograde tracing using the cholerae-toxin B (CTB) tracer. CTB tracer conjugated with Alexa Fluor-555 (red fluorescence) was injected into the PnC (Fig. 1a), revealing CTB-labelled neurons in prefrontal regions (Fig. 1b). The majority of labelled cells (92.78 %, 1053 of 1135 neurons; n = 3 mice) were found in layer V of the dorsal anterior cingulate cortex (dACC) and the secondary motor cortex (M2; Fig. 1c), whereas only 6.96 % and 0.26 % were found in other prefrontal sub-regions – namely the ventral anterior cingulate cortex (vACC; with prelimbic cortex, PL) and the infralimbic cortex (IL), respectively. Hereafter, we refer to these neurons as the dACC→PnC neurons.

**Figure 1.**
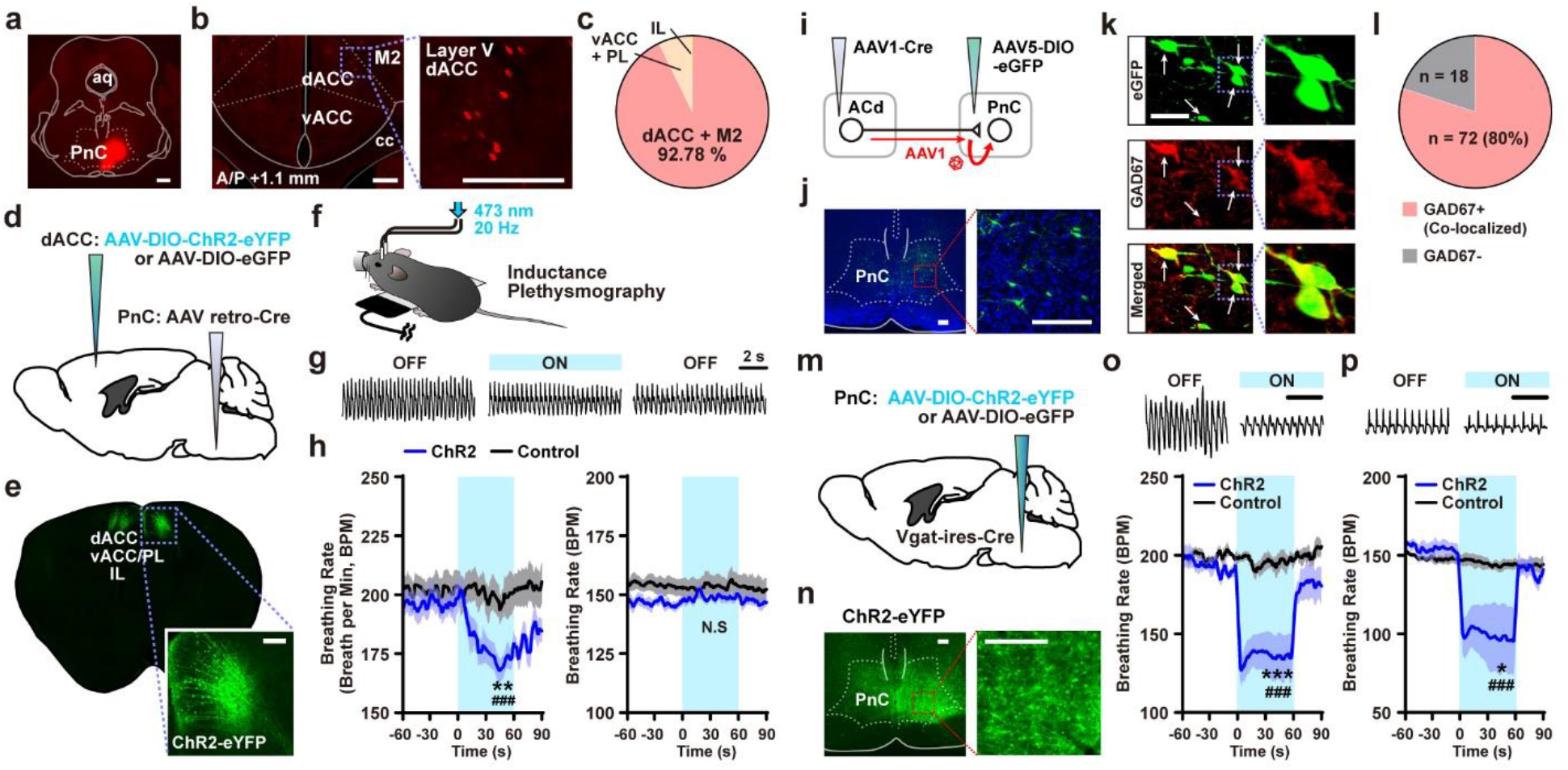
Identification of a top-down slow breathing circuit. **a**, Injection of CTB-Alexa Fluor 555 tracer into the PnC. **b**, CTB-labelled neurons in the dACC and M2. **c**, Quantification of CTB-labelled neurons in prefrontal subregions. **d**, Schematic showing projection-specific expression of AAV constructs in dACC→PnC neurons. **e**, ChR2-eYFP expression in dACC→PnC neurons. **f**, Inductance plethysmography experiment with photostimulation under light anesthesia. **g**, Raw breathing traces show changes in breathing induced by photostimulation in the ChR2 group. Scale bar, 2 s. **h**, Photoactivation of dACC→PnC neurons led to a decrease in breathing rate from a baseline of 200 BPM (left), but not from a baseline of 150 BPM (right; n = 6 in ChR2, n = 5 in eGFP). **i**, Schematic of anterograde neuronal labelling with AAV1-Cre injection. **j**, Downstream PnC neurons expressing DIO-eGFP. **k**, Confocal images showing the co-localization of eGFP-labelled neurons and GAD67. **l**, Quantification of co-localized neurons. **m**, Schematic of Cre-dependent AAV expression in PnC^GABA^ neurons. **n**, ChR2-eYFP expression in PnC^GABA^ neurons. **o**, **p**, Photoactivation of PnC^GABA^ neurons decreased breathing rates from both 200 BPM (**o**) and 150 BPM baselines (**p**). Top, Raw breathing signals. Scale bar, 2s. Bottom, breathing rates shown in group average (n = 5 ChR2, n = 4 control). aq, aqueduct. PnC, pontine reticular nucleus caudalis. dACC, dorsal anterior cingulate cortex. vACC, ventral anterior cingulate cortex. PL, prelimgic cortex. IL, infralimbic cortex. Scale bar, 500 μm in (**a**), 200 μm in **b**, **e**, **j**, **n**, 50 μm in **k**. **h**, **o**, **p**, Repeated measures (RM) two-way ANOVA followed by Sidak’s post-hoc test. ^N.S^P > 0.05, *P < 0.05, **P < 0.01, ***P < 0.001 between-group comparison; #P < 0.05, ##P < 0.001 within-group comparison. Data are shown as mean ± s.e.m.

To investigate the role of dACC→PnC neurons in breathing control, we used a projection-specific labelling strategy to express the light-driven actuator, Channelrhodopsin-2 (ChR2), in the dACC. A retrograde adeno-associated viral vector encoding Cre recombinase (AAVretro-Cre) was unilaterally injected into the PnC. Another AAV vector that expresses ChR2 (fused with eYFP; AAV-DIO-ChR2-eYFP; n = 6) or green fluorescence protein (for control group; AAV-DIO-eGFP; n = 5) in the presence of Cre was bilaterally injected into the dACC (Fig. 1d). This resulted in the successful expression of eYFP or ChR2-eYFP in the dACC (Fig. 1e). Consistent with our tracing results (Fig. 1c), only layer V neurons in the dACC/M2 regions were labelled with eYFP. To eliminate confounding effects of locomotion on breathing, these mice were carefully anesthetized and placed in a stereotaxic frame, and breathing was monitored using inductance plethysmography (Fig. 1f). Baseline breathing rate under light anesthesia was ~200 breaths per minute (BPM). Notably, photoactivation of the dACC→PnC neurons significantly reduced the breathing rate in the ChR2 group to below 170 BPM on average (Fig. 1g and 1h left). No change in breathing rate was observed for controls (Fig. 1h left). Based on these results, we hypothesized that dACC→PnC neurons are cortical breathing modulators that induce slow breathing upon activation.

The same mice were also tested in a slightly deeper anesthetic state, where the baseline breathing rate was 150 BPM. Interestingly, no change in breathing rate was observed in both the ChR2 and control groups in response to dACC→PnC neuronal activation (Fig. 1 h right), suggesting that this cortical breathing circuit, unlike pontomedullary breathing centers, cannot override homeostatic respiratory controls and may function only above a certain homeostatic threshold.

Based on our search of the Allen Mouse Brain Atlas, we predicted that dACC neurons target GABAergic inhibitory neurons in the PnC (Extended data Fig. 1). To test this idea, we performed anterograde tracing using serotype 1-packaged AAV (AAV1), which travels to efferent (downstream) neurons in a trans-synaptic manner^16,17^. An AAV1 vector encoding Cre recombinase (AAV1-Cre) was injected into the dACC (Fig. 1i) and AAV-DIO-eGFP was injected into the PnC. The result was sparse but successful expression of eGFP in PnC neurons (Fig. 1j; 90 cells in 3 mice). To determine which types of cells were labelled, we immunolabelled PnC coronal sections with antibodies against GAD67, a marker of GABAergic cells. In total, 72 of 90 cells (80%) were co-labelled with eGFP and GAD67 (Fig. 1k and 1l), indicating that dACC neurons project to inhibitory neurons in the PnC.

We next tested the role of PnC GABAergic neurons (PnC^GABA^) in breathing control using optogenetics. We injected AAV-DIO-ChR2-eYFP (Fig. 1m and 1n), AAV-DIO-eNpHR3.0 (an AAV that expresses halorhodopsin-3.0 in the presence of Cre; Extended data Fig. 2a), or control AAV-DIO-eGFP into the PnC of *Vgat-ires-Cre* transgenic mice, which express Cre in GABAergic neurons. During testing of lightly anesthetized mice, breathing rhythms were monitored using inductance plethysmography (baselines of 200 or 150 BPM). When photostimulation was applied, a dramatic decrease in breathing rate was observed in the ChR2 group for both breathing baseline conditions (Fig. 1o and 1p), whereas the control group exhibited no change. Conversely, an increase in breathing rate was observed when PnC^GABA^ neurons were photoinhibited via eNpHR3.0 (Extended data Fig. 2a-d). These data indicate that PnC^GABA^ neurons are inhibitory regulators of breathing rhythm, and their activation slows breathing. Therefore, inputs from the prefrontal area (dACC/M2) to the PnC comprise a top-down modulatory circuit that induces slow breathing.

### Neuronal activity correlates with breathing changes during purposeful behaviors

Breathing can be profoundly controlled by behavior. In particular, behaviors that require oropharyngeal activities and airway protection can perturb or entrain breathing cycles^1,5,7^. We hypothesized that dACC→PnC neurons and their downstream PnC^GABA^ targets respond to purposeful behaviors that require synchronization of breathing, such as drinking. While coordination of breathing and swallowing behavior is generally controlled by trigeminal reflexes^18^, accumulating evidence suggests that apnea during drinking also involves inputs from high-order cortices^14,15,19^. For example, distinct breathing patterns are seen during conscious drinking and unconscious swallowing^14^, and neuroimaging studies have revealed that the dACC and supplementary motor area (SMA) are activated during the oral/preparatory phase before swallowing^15,19^.

To monitor breathing signals in freely behaving mice, we implanted a micro-thermistor sensor into the nasal cavity^2,7,20,21^ (Fig. 2a). Temperature difference between inspiratory and expiratory breathing cycles was converted into voltage signals by the sensor. To simultaneously monitor breathing rhythms and dACC→PnC neuronal activity (Fig. 2a and 2c), the same group of mice received another round of surgery to express the calcium-activity indicator jGCaMP7s. Similar to Fig 1d, AAVretro-Cre vector was injected into the PnC, and another AAV carrying a Cre-dependent jGCaMP7s cassette (AAV-FLEX-jGCaMP7s) was unilaterally injected into the dACC (Fig. 2c). We monitored breathing and calcium signals in mice as they voluntarily drank water (Fig. 2b). dACC→PnC neurons were transiently activated upon the oral phase of drinking behavior, which was defined as the moment a bolus of water was taken into the mouth cavity (Fig. 2d and 2e). Downstream PnC^GABA^ neuronal activity was monitored using transgenic mice (*Vgat-ires-Cre*) that express the jGCaMP7s indicator in the PnC (Fig. 2f). Similarly, prominent increases in PnC^GABA^ neuronal activity were observed during drinking events (Fig. 2g and 2h). Therefore, consistent with previous human studies, our results demonstrate that a top-down breathing circuit and its downstream targets are activated during drinking behaviors to coordinate breathing rhythms.

**Figure 2.**
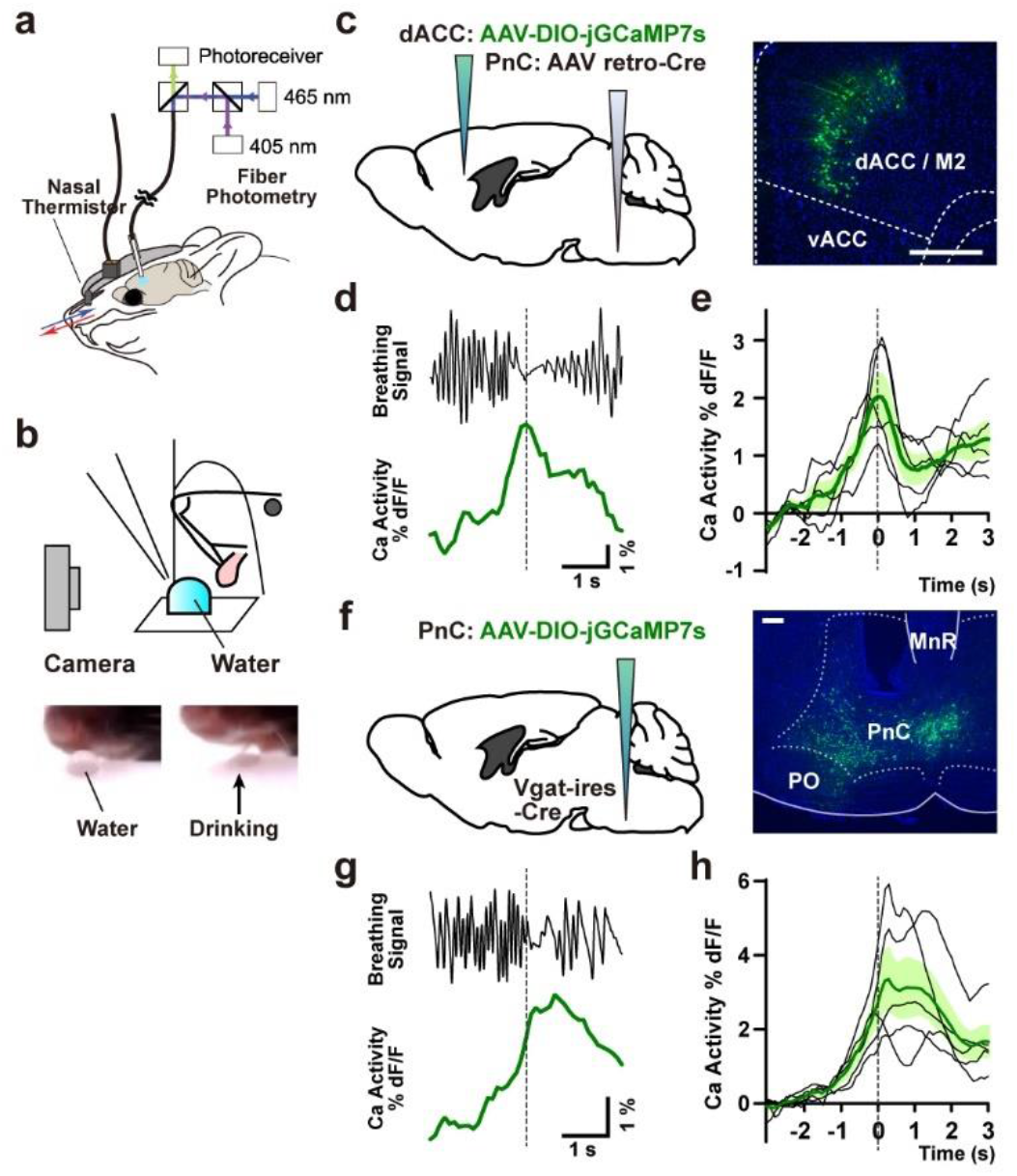
dACC→PnC and PnC^GABA^ neuronal activities are correlated with drinking behavior and breathing rhythms. **a**, Simultaneous recording of breathing cycles and neuronal calcium levels. **b**, Schematic showing the observation of drinking behavior. **c**, Surgical procedure for expressing the jGCaMP7s indicator in dACC→PnC neurons (left), and image showing jGCaMP7s expression (right). **d**, **e**, Representative traces showing breathing cycles, dACC→PnC activity (**d**), and average dACC→PnC activity (**e**; n = 5 mice) during drinking behavior. **f**, Schematic of the expression of jGCaMP7s in PnC^GABA^ neurons (left), and image showing jGCaMP7s expression (right). **g**, **h**, Representative traces showing breathing, PnC^GABA^ activity (**g**), and average PnC^GABA^ activity (**h**; n = 5 mice) during drinking behavior. Scale bar, 200 μm. Data are shown as mean ± s.e.m.

We next monitored breathing rhythms and dACC→PnC or PnC^GABA^ neuronal activity during other behaviors that involve oropharyngeal activities, namely swimming, squeaking (vocalization), and sniffing (Extended data Fig. 3). Two types of swimming tests were performed. Mice were briefly submerged in water (1–2) and then either immediately removed from the water (withdrawal test) or allowed to swim after release. During the withdrawal test (Extended data Fig. 3a-e), apneic breathing cycles were observed only during submersion and fast breathing cycles resumed after withdrawal. Consistent with these breathing changes, short (~2 s) increases in dACC→PnC and PnC^GABA^ neuronal activity were induced during submersion and decreases in neuronal activity were observed after withdrawal. The release/swim test resulted in dramatic changes in breathing patterns with repeated apneic cycles (Extended data Fig. 3f-j). Both dACC→PnC and PnC^GABA^ neurons showed sustained increases in calcium signals, revealing a correlation between neuronal activity and slow-apneic breathing cycles. While the extent to which the cerebral cortex modulates breathing rhythms during swimming is not yet fully understood (due to technical limitations), evidence from humans and rats suggests that the activation of prefrontal and motor cortical regions may be involved in controlling breathing during swimming. Our data are consistent with top-down pathways helping to modulate breathing rhythms during swimming, working in combination with the well-established trigeminal diving reflex.

**Figure 3.**
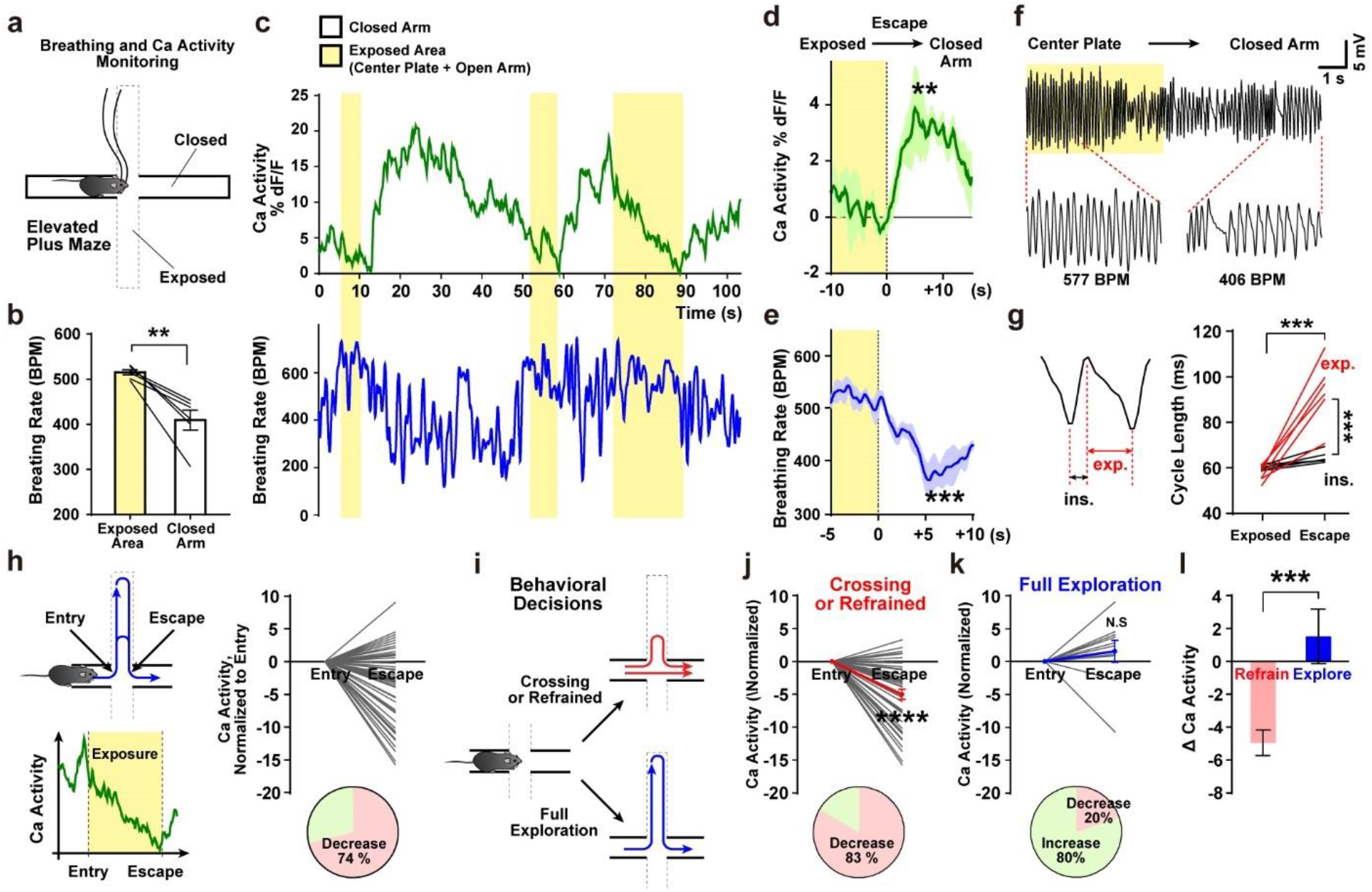
dACC→PnC neuronal activity is correlated with breathing and anxiety-related behaviors. **a**, Simultaneous monitoring of breathing cycles and calcium activity during EPM testing. **b**, Breathing rate was lower in the closed arms compared to exposed areas (n = 6). **c**, Representative traces of calcium levels and breathing rate during EPM testing. **d**, dACC→PnC neuronal activity increased during escape events (exposed → closed arm). **e**, Breathing rate decreased during escape events. **f**, Raw breathing trace showing the changes in breathing cycles after escape. **g**, Analyzing the length of inspiratory and expiratory phases before and after escape. **h**, Normalized change in dACC→PnC activity during exploration of exposed areas. **i**, Classification of behavioral decisions. **j-l**. Normalized activity changes in refrained (**j**) or full exploration (**k**) episodes. Changes in calcium levels were observed in classified episodes (**l**). **b**, **d**, **e**, **k**, two-tailed paired t test; **g**, RM two-way ANOVA followed by Sidak’s post-hoc test; **j**, Wilcoxon signed rank test; **k**, paired t test; **l**, Mann-Whitney U test. **P < 0.01, ***P < 0.001, and ****P < 0.0001. Data are shown as mean ± s.e.m.

Finally, we monitored the activity of dACC→PnC neurons during foot shock-induced vocalization or sniffing. Shock-induced vocalizations (squeaks) were only seen during slow expiratory breathing cycles, resulting in a decrease in breathing rate. Notably, these vocalizations were associated with an increase in dACC→PnC activity, again revealing a correlation between activation of these neurons and slow breathing rate (Extended data Fig. 3k-n). In contrast to drinking, swimming, and squeaking, which require slow or apneic breathing cycles, sniffing is a behavior that results in faster breathing cycles (as much as 600 BPM). Following contact with an object and the onset of sniffing, decreased dACC→PnC neuronal activity was observed. Rebound-like activation was then observed upon the termination of sniffing (Extended data Fig. 3o-r). These data suggest that dACC→PnC neurons receive inhibitory inputs when the animal is engaged in fast-breathing behavior (Extended data Fig. 3o-r).

Taken together, data acquired using the nasally implanted thermistor sensor revealed that activity patterns of the dACC→PnC pathway are tightly correlated with changes in breathing patterns during behaviors that require coordinated changes in breathing. While many of these behaviors are controlled by somatic and trigeminal reflexes, our findings suggest that a top-down, slow-breathing pathway is also involved in regulating the behavioral entrainment of breathing, presumably operating at different stages of the entrainment process.

### dACC→PnC activity correlates with emotionally evoked breathing changes

Hyperventilation and shortness of breath are prominent symptoms of anxiety and panic disorders in humans^3,22,23^, whereas long, slow breaths or mindfulness skills can help alleviate anxiety^1,11,24^. We next investigated how dACC→PnC neurons respond to anxiety-provoking environments and how these responses correlate with behavior and breathing patterns. We recorded calcium responses and breathing (as in Fig. 2a) while mice were placed in an elevated plus maze (EPM; Fig. 3a). All mice tested (n = 6) exhibited an increase in breathing rate while in exposed areas (center or open arms) compared to when they were in closed arms (Fig. 3b). Consistent with our previous study^2^, we found a precise correlation between the location of the mouse and its breathing rate (Fig. 3c-f). We observed slow breathing cycles after “escape” events – when a mouse moved from an exposed area to a closed arm (Fig. 3e and 3f). Increases in dACC→PnC activity were also observed during these escape events (Fig. 3d), revealing a correlation between slow breathing cycles and increased dACC→PnC neuronal activity. A detailed analysis of breathing phases further revealed that elongation of the expiratory phase (post-inspiration through expiration) contributed to the slower breathing rate (Fig. 3g).

We further analyzed dACC→PnC activity patterns during the voluntary exploration of open arms. During each episode, the change in dACC→PnC neuronal activity was calculated by subtracting the calcium level at open-arm entry from the calcium level at open-arm exit (Fig. 3h). This revealed an interesting correlation between neuronal activity and mouse behavior in the open arms. Prominent decreases in dACC→PnC activity were observed when mice failed to reach the end of the open arm, but instead quickly returned to the center (refrained behavior, Fig. 3j). By contrast, neuronal activity persisted or gradually increased in most episodes (80%) in which the mouse fully explored the open arm (full exploration, Fig. 3k and 3l). These analyses reveal a correlation between the response of dACC→PnC neurons and anxiety-related behaviors. In another set of experiments, mice were exposed to a similar but inescapable anxiogenic environment (Extended data Fig. 4). For these experiments, mice were placed in a closed cylinder on an elevated platform and then the cylinder was removed to expose them to height. This exposure elicited a steep increase in breathing rate (reaching ~600 BPM) and a concurrent decrease in dACC→PnC activity (Extended data Fig. 4).

Taken together, these experiments show that dACC→PnC neuronal activity is correlated with breathing and behavior in anxiety-provoking environments. Specifically, we observed increases in dACC→PnC neuronal activity during escape events, which were coupled with slow breathing cycles characterized by longer exhales. dACC→PnC activity decreased in anxiogenic environments (open arms of an elevated plus maze or an elevated platform). However, persistent dACC→PnC activity was observed when mice overcame an anxiogenic condition and fully explored the environment, suggesting that increased dACC→PnC activity promotes relief from anxiety.

### The dACC→PnC circuit alleviates anxiety-like behaviors

Our calcium imaging results revealed that dACC→PnC neuronal activity was inhibited by anxiety-provoking environments, whereas neuronal activity persisted when mice overcame an anxiogenic condition. We therefore hypothesized that artificial activation of dACC→PnC neurons would suppress anxiety-like behaviors in mice and drive exploratory behaviors. To selectively activate dACC→PnC neurons in anxiety tests, we used the same viral strategy as in Fig. 1d-e to bilaterally express ChR2 in dACC→PnC neurons (Fig. 4a). We first performed a control test using a real-time place preference (RTPP) paradigm, which does not promote anxiety-like behaviors but instead reveals potential changes in valence (see the Methods section for details regarding the RTPP paradigm). During the second epoch of testing (minute 10–20 of a 30-min session), light stimulation was applied when the mouse was in one chamber of a two-chamber box. Following this stimulation, neither the ChR2 nor control groups displayed preference or aversion for the light-paired chamber (Fig. 4b), indicating that dACC→PnC activation does not alter valence. As a second control test, one chamber of the two-chamber box was paired with the odor of a female mouse cage to provoke male-approach behavior. Male mice showed strong preference for the odor-paired chamber, and this behavior was not altered by photoactivation (Fig. 4c). Next, these mice were tested in an aversive environment where one side of a cage was paired with fox anogenital odor – trimethylthiazoline (TMT)^25^. Mice avoided the TMT odor, but this avoidance was abolished by dACC→PnC photostimulation in the ChR2 group but not in controls (Fig. 4d). Similar results were seen for other tests that probe behaviors associated with fear and anxiety, such as the elevated plus maze test (Fig. 4e), the looming-hiding task^26^, and the light-dark choice test (Extended data Fig. 5). Photostimulation of the ChR2 group increased exploration of the open arms in the EPM test (Fig. 4e) and exposed regions of each test arena. Taken together, these results show that activation of dACC→PnC neurons consistently alleviates anxiety-like responses to aversive stimuli without altering valence or approach behaviors.

**Figure 4.**
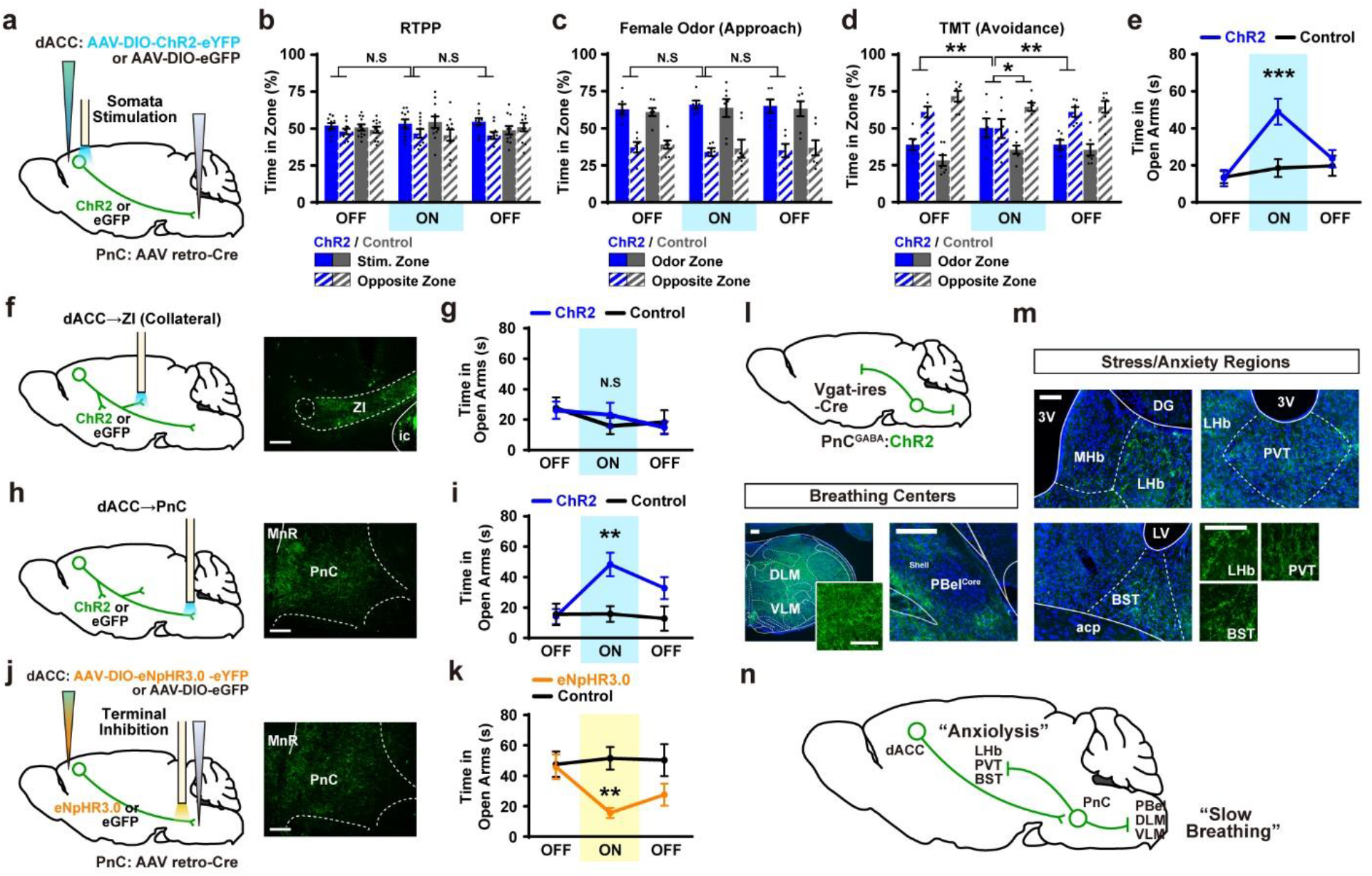
dACC→PnC neurons and downstream circuit alleviate anxiety-like behaviors. **a,** Schematic showing photoactivation of dACC→PnC neurons (somata) expressing ChR2-eYFP. **b-d**, Behaviors in the RTPP test (**b**; ChR2, n = 10; eGFP, n = 11) and approach response to female odor (**c**; ChR2, n = 6; eGFP, n = 7) were not altered by photoactivation. Avoidance response to TMT was reduced by photoactivation of dACC→PnC neurons (**d**; ChR2, n = 6; eGFP n = 7). **e**, Photoactivation of dACC→PnC neurons increased open arm exploration in the EPM test (ChR2, n = 10; eGFP, n = 11). **f**, **g**, Photoactivation targeting dACC→ZI collaterals (**f**) did not affect open arm exploration during EPM test (**g**, ChR2, n = 8; eGFP, n = 7). **h**, **i**, Photoactivation targeting dACC→PnC projection terminals (**h**) increased open arm exploration during EPM test (**i**, ChR2, n = 7; eGFP, n = 7). **j**, **k**, Photoinhibition targeting dACC→PnC projection terminals (**j**) reduced open arm exploration during EPM test (**k**, eNpHR3.0, n = 10; eGFP, n = 11). Images (**f**, **h**, **j**) show the histology of projection terminals labelled with eYFP. **l**, **m**, Terminals of PnC^GABA^ neurons expressing ChR2-eYFP. eYFP-labelled axons were observed in pontomedullary breathing centers – DLM, VLM, and PBel^Shell^ (**l**) and stress/anxiety-related forebrain regions – LHb, PVT, and BST (**m**). **n**, Divergent connectivity of PnC neurons target breathing centers and fear/anxiety-related regions. ZI, zona incerta; ic, internal capsule; MnR, median raphe nucleus; DLM, dorsolateral medulla; VLM, ventrolateral medulla; PBel, exterior-lateral parabrachial nucleus; MHb, medial habenula; LHb, lateral habenula; DG, dentate gyrus; PVT, paraventricular nucleus of thalamus; 3V, third ventricle; BST, bed nucleus of stria terminalis; LV, lateral ventricule; acp, anterior commissure, posterior. Scale bar, 200 μm in **f**, **h**, **j**, **l**, and 100 μm in **m**. RM two-way ANOVA followed by Sidak’s post hoc test (**b**-**e**, **g**, **i**, **k**). *P < 0.05, **P < 0.01, ***P < 0.001. Data are shown as mean ± s.e.m.

We then sought to characterize the circuit that acts downstream of dACC→PnC neurons to promote anxiolysis. We reasoned that dACC→PnC projections could drive changes in both breathing rate and emotion. Alternatively, dACC→PnC projections could control breathing, whereas axon collaterals of these neurons could target other downstream regions to alleviate negative affect. We found that dACC→PnC neurons send putative collaterals to several other regions, including the dorsal striatum, the mediodorsal and ventral midline thalamic nuclei, the zona incerta, and the deep mesencephalic nucleus of the midbrain (Extended data Fig. 6). Using mice that express ChR2-eYFP in dACC→PnC neurons, we photostimulated axon collaterals within the zona incerta (ZI; Fig. 4f), a brain region known to regulate fear- and anxiety-related responses in mice^27,28^. This did not affect anxiety-like behavior in the elevated plus maze test (Fig. 4g). By contrast, stimulating terminals within the PnC increased the time spent exploring the open arm (Fig. 4h and 4i). Using mice that express eNpHR3.0-eYFP in dACC→PnC neurons, we asked whether inhibiting dACC→PnC projections affected anxiety-like behavior (Fig. 4j). No changes in behaviors were observed in control tests, namely the real-time place aversion or female odor preference tests (Extended data Fig. 7). In the EPM test, however, photoinhibition of dACC→PnC terminals reduced time spent exploring the open arm (Fig. 4k). Taken together, these results indicate that dACC→PnC projections are important for controlling anxiety-related responses in threatening environments.

Given these results, we next characterized the downstream projections of PnC^GABA^ neurons. Using *Vgat-ires-Cre* transgenic mice that express ChR2-eYFP in the PnC, we found eYFP-labelled axons in a wide range of medullary structures (Fig. 4l) covering the dorsolateral and ventrolateral medulla (DLM and VLM), as well as the shell of the external parabrachial nucleus (PBel^Shell^) in the pons, regions critically involved in pneumotaxic and rhythmogenic control of breathing^2,29^. Importantly, eYFP-labelled fibers were also found in a few forebrain regions (Fig. 4m) including the lateral habenula (LHb), the paraventricular thalamic nucleus (PVT), and the bed nucleus of the stria terminalis (BST), structures implicated in fear and anxiety responses^30–33^. Therefore, the practice of slow breathing may alleviate anxiety and negative affect in parallel through divergent connectivity of the PnC^GABA^ neurons (Fig. 4n).

### The dACC→PnC circuit is required to coordinate breathing and behavior

Throughout the study, we observed activation of dACC→PnC neurons during behaviors that must be coordinated with slow breathing rhythms. To determine whether dACC→PnC neurons coordinate breathing and behavior, we inhibited the dACC→PnC circuit and assessed the effect on both breathing cycles and associated behaviors, namely drinking and anxiety-related behaviors. To photoinhibit projection terminals, surgical procedures were performed to express eNpHR3.0-eYFP (or control eGFP) in dACC→PnC neurons (Fig. 5a). Fiber optic elements were implanted into the PnC region to deliver light stimuli, and thermistor sensors were implanted in the nasal cavity to simultaneously record breathing rhythms.

**Figure 5.**
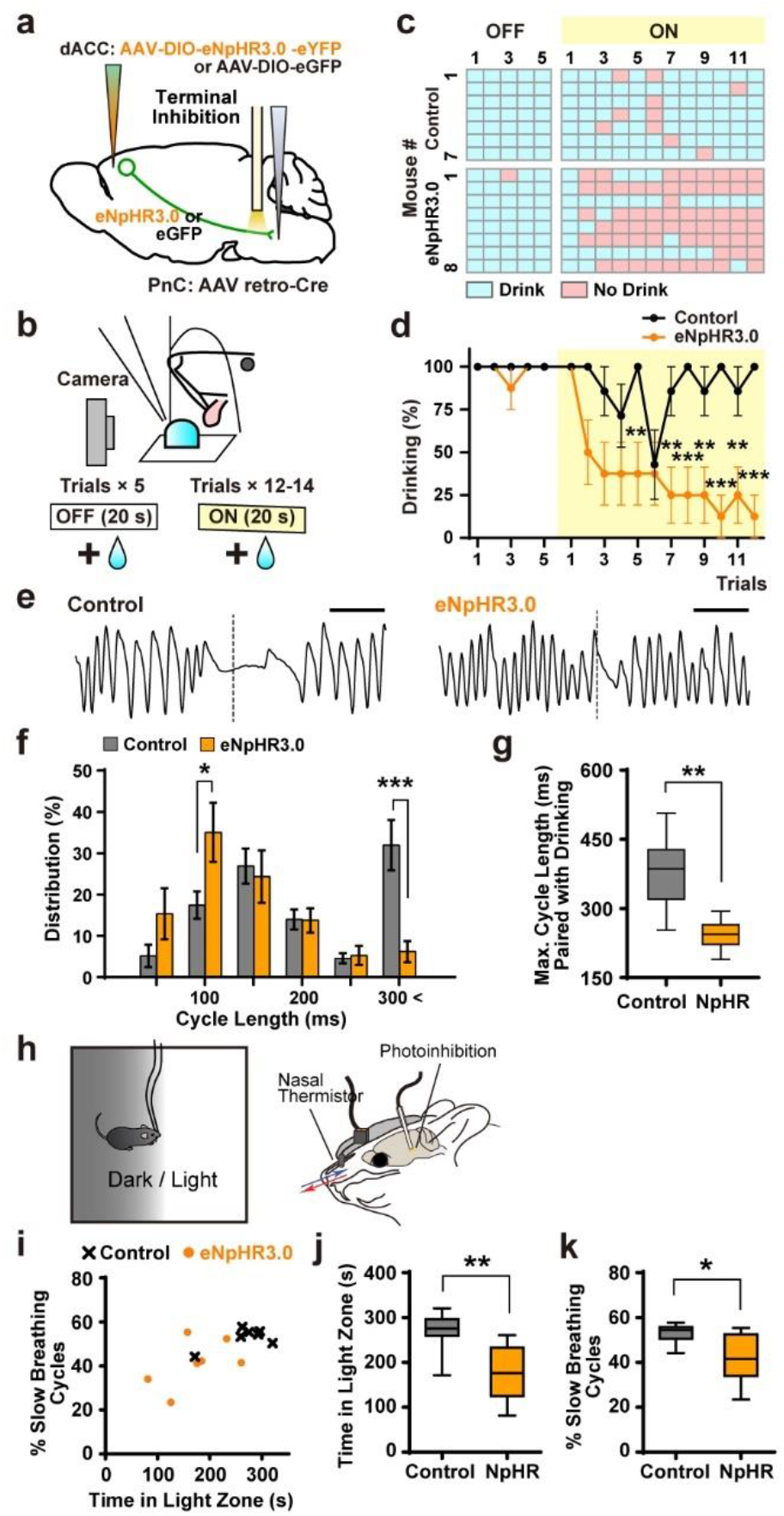
dACC→PnC terminal inputs are required for behaviors associated with slow breathing. **a**, Schematic of the photoinhibition of dACC→PnC terminals **b**, Schematic showing the observation of drinking behavior. **c**, **d**, Successful drinking responses are shown as individual data (**c**) or average traces (**d**) (eNpHR3.0, n = 8; eGFP, n = 7). **e**, Representative traces showing breathing cycles paired with drinking in control (eGFP) or eNpHR3.0-expressing mice. Scale bar, 500 ms. **f**, Percent distribution of the length of breathing cycles seen during drinking. **g**, Average of maximal cycle lengths paired with drinking episodes (eNpHR3.0, n = 8; eGFP, n = 7). **h**, Schematic of light/dark choice test with breathing monitoring and photoinhibition. **i-k**, Breathing (**i**, **k**, % of breathing cycles slower than the median rate) and behavioral correlates (**i**, **j**, time in light zone) observed in the light/dark choice test (eNpHR3.0, n = 7; eGFP, n = 7). RM two-way ANOVA followed by Sidak’s post hoc test (**d**, **f**, **g**, **j**, **k**). *P < 0.05, **P < 0.01, ***P < 0.001. Bar graphs are shown as mean ± s.e.m. Box-whisker plots are shown as median and interquartile range with 5-95 percentile distribution.

We first inhibited the dACC→PnC circuit and assessed the impact on voluntary drinking and paired breathing cycles. After overnight dehydration, mice in both groups (eNpHR3.0-eYFP or control eGFP) drank successfully (i.e., took the water bolus into their mouth) when water droplets were presented for 20 s without light stimulation. On the first drinking episode during the light-ON session, both groups drank successfully, but in subsequent episodes the eNpHR3.0 group exhibited refusal behaviors and decreased drinking success compared to the control group (Fig. 5c and 5d). Once trials were no longer paired with photoinhibition, drinking success rates for the eNpHR3.0 group recovered to > 70% by the third session (Extended data Fig. 8). Drinking responses of the control group were highly successful (> 80%) throughout the testing session. To determine whether breathing was affected by photoinhibition, we analyzed breathing cycle lengths during drinking behavior (defined as −100 to +500 ms from drinking; Fig 5e). Compared to the control group, eNpHR3.0 mice exhibited shorter breathing cycles during drinking behavior (fewer breathing cycles that exceeded 300 ms and more breathing cycles of 100-150 ms) (Fig. 5f). Importantly, the eNpHR3.0 group paired shorter apneic cycles with drinking episodes than controls (Fig 5g). These data indicate that the dACC→PnC circuit is required to successfully coordinate slow breathing cycles and voluntary drinking behavior. Thus, the trigeminal pathway mediates reflex responses during unconscious swallowing, whereas top-down pathways may play a critical role in the preparatory control of breathing cycles to ensure safe coordination.

Finally, to explore the effect of dACC→PnC inhibition on anxiety-related behaviors, we utilized a light/dark choice model that probes anxiety-related states based on exploration of a light zone. We designed a light/dark testing chamber in which half of an open square arena was shaded (Fig. 5h). Mice could move freely between light and dark zones with minimal hinderance from patch cords used to monitor breathing and deliver light. While in the light zone, mice received 590-nm light stimuli to inhibit dACC→PnC terminals. Notably, mice in the eNpHR3.0 group spent less time in the light zone compared to controls (Fig. 5i and 5j). Consistent with this behavioral change, the eNpHR3.0 group exhibited a smaller proportion of slow breathing cycles (cycles slower than the median breathing rate throughout testing) when in the light zone (Fig. 5k), indicating that the dACC→PnC circuit is required to induce slow breathing cycles to relieve anxiety. Taken together, these analyses demonstrate that the dACC→PnC circuit is necessary to coordinate behaviors with slow breathing.

## Discussion

Most breathing research has targeted brainstem networks that receive somatic input to maintain gas homeostasis (pH, blood O2, and CO2 levels) and to generate rhythmic breath cycles through cranial and spinal motor pathways^1,5^. Compared to these well-characterized networks in the deep brainstem, how cortical circuits and their projections modulate breathing rhythms remains poorly understood. Here we describe a prefrontal circuit that projects to inhibitory neurons in the pons, which in turn project widely to breathing centers in the pons and medulla. The dACC→PnC circuit, along with downstream PnC^GABA^ neurons, promote slow breathing cycles and respond to behaviors such as drinking to entrain slow/apneic breathing cycles. Taken together, here we report the discovery of a top-down brain circuit for the inhibitory control of breathing and the coordination of breathing and behavior.

It has been proposed that neurons in the primary motor cortex that bypass pontomedullary breathing circuits and project directly to the spinal cord (the corticospinal tract) are crucial for the volitional/behavioral control of breathing^12^. However, this theory has limitations, as it is unclear how these cortical circuits would incorporate on-going functions of autonomous breathing circuits instead of interfering with them^4^. Moreover, accumulating evidence in mice and rats indicates that medullary and pontine neuronal responses are correlated with and required for the entrainment of breathing rhythms to sniffing behavior^1,2,7,34^. Multiple studies in a wide range of species (including cats, birds, and monkeys) report that the parabrachial nucleus of the pons plays a critical role in coordinating vocalizations and respiratory rhythms^35–37^. These behaviors are used to voluntary interact with objects, environments, and conspecifics, and thus must be tightly synchronized with behavioral status^1,38^. Therefore, it is unlikely that behavioral inputs simply bypass the pontomedullary networks. Instead, pontomedullary breathing networks are likely modulated by top-down inputs.

Various voluntary efforts to change breathing patterns (e.g., by emphasizing longer exhales or box breathing) are used to control emotions^24,39–42^. However, it is not fully understood how these practices modulate the internal state of the brain. One line of research explores changes in bodily states and feedback interoception. The thinking is that longer expiration (exhale) leads to a decreased heart rate via respiratory sinus arrhythmia^43^, which in turn affects the interoceptive perception of the heart rate^44^. Another line of research explores the role of medullary breathing centers themselves. Neurons in the preBötzinger complex that express Cdh9 and Dbx1 respond to the inspiratory phase of breathing in mice^45^. These neurons project to noradrenergic neurons in the locus coeruleus. Therefore, fast or slow breathing rhythms may modulate the activity of noradrenergic neurons, which then influence the brain-wide state of arousal. Along with these bottom-up mechanisms, our current findings reveal a complementary mechanism in which a top-down breathing circuit is critical for controlling negative emotions. Inhibitory PnC neurons, which receive input from the anterior cingulate cortex, project to medullary breathing centers but also to anxiety-related regions in the forebrain. Given this connectivity, the alleviation of negative affect may occur simultaneously with volitional efforts to slow breathing cycles.

## Data availability

All data are available from the corresponding author upon request.

## Acknowledgments

We thank all members of the Han laboratory for scientific discussions and support. J.J. is supported by IRGS 2020-1710 from the KAVLI Institute for Brain and Mind (KIBM). S.H. is supported by 5R01MH116203 from NIMH.

## Author contributions

S.H. and J.J. designed the study and secured funding. J.J., D.D.O., and S.H. wrote the manuscript. J.J. performed the experiments and analyzed the data. S.L. helped with thermistor implantation surgeries.

## Methods

### Mice

All procedures for animal surgical and behavioral experiments were approved by the institutional animal care and use committee (IACUC) of the Salk Institute according to NIH guidelines for animal experimentation. Wildtype or *Vgat-ires-Cre* transgenic lines with C57Bl/6 background were group-housed on a 12-h light/dark cycle under constant temperature and humidity conditions, and provided with food and water ad libitum, except experiments with single housing or overnight restriction of water.

### Respiratory measurements

#### Inductance plethysmography

Pressure signals generated by respiratory chest movements were converted into voltage signals by a piezoelectric sensor placed beneath the chest of mice under isoflurane-induced anesthesia. To induce anesthetic conditions with varying breathing rates, the duration and concentration of isoflurane treatment were carefully adjusted. Mice were initially exposed to 3% isoflurane. Five-ten seconds after the loss of movement, mice were moved to a stereotaxic frame. Isoflurane concentration (delivered through the nasal cone of the stereotaxic device) was adjusted to 0.8-1% or 1.2-1.5%, to induce light (with ~200 breathing cycles per minute; BPM) or deep (~150 or 100 BPM) anesthetic conditions. The piezoelectric sensor was connected to a PowerLab monitoring system which was operated by LabChart 8 Pro software (ADInstruments). LabChart 8 Pro software was used for the sampling of voltage signals, signal processing, detection of breathing cycles, and the calculation of breathing rate or amplitude. Data were sampled at 400 Hz, low-pass filtered at 10 Hz, and smoothed with a 100-ms window. (Fig. 1h, 1o, and 1p). For statistical analysis, the 150-second data of breathing rate was divided into 30-second periods and averaged within each period. The plots (Fig. 1h, 1p and 1q) represent 150 bins (1-s each) of the raw data obtained with 400-Hz sampling rate.

#### Breathing monitoring with nasal thermistor sensor

Nasal thermistor-based breathing monitoring was performed as described^2,46^. The custom sensor was built with an NTC thermistor (TE Connectivity) and an interconnector (Mill-Max), and stereotaxically implanted in the nasal cavity of mice, with approximately 1 mm of depth from the skull surface. During the monitoring experiments in awake behaving mice, the interconnector was attached to an electric patch cord, which then connects to a rotary joint with a voltage divider (Phidgets) and to the PowerLab device. Temperature difference between inspiratory and expiratory airflow was converted into voltage signals. Data were sampled at 1 kHz, filtered with a 0.4-25 Hz band-pass filter, and smoothed with 50-ms moving window. Breathing cycles, rate, and estimated amplitude were automatically calculated by the LabChart Pro 8 software, and peak detection was validated with manual observation.

### Stereotaxic surgery

Mice were anesthetized with isoflurane (4% initial, 1.5% for maintenance) and fixed on a stereotaxic surgery frame (David Kopf Instruments) equipped with a heating pad. The skull was exposed using scalpels, and the cranium was drilled with a handpiece drill (Foredom). Viral solutions or the CTB tracer were loaded into glass pipettes filled with mineral oil, then injected into target brain coordinates at the rate of 1 nL per second using a Nanoject III programmable injector (Drummond Scientific). Target brain coordinates (relative to bregma) for virus and CTB tracer injections were: anterior-posterior (AP) +1.2 mm, medial-lateral (ML) ±0.5 mm, dorsal-ventral (DV) −1.2 mm for the bilateral dACC, and +15 degrees angled, AP −5.0 mm, ML +1.8 mm, DV −5.4 mm, for the unilateral PnC. Behavioral experiments with optogenetics or calcium imaging were performed 3-4 weeks after the injection surgery. Nasal thermistors were implanted on the same day of injection surgeries. In case thermistors were implanted alone (without virus injection), mice were allowed to recover for 7 days after the surgery.

For optogenetic control of dACC→PnC neurons or projection terminals, 400-nL solution of AAVdj-EF1a-DIO-hChR2-eYFP-WPRE-pA (1.2E+12 GC/mL) or AAV5-EF1a-DIO-eNpHR3.0-eYFP-WPRE-pA (6.1E+12 GC/mL GC/mL) or control AAVdj-syn-DIO-eGFP (2.3E+12 GC/mL) was injected into the dACC coordinate, and 200-nL solution of AAV retrograde-EF1a-Cre-WPRE-pA (GC/mL) was injected in the PnC. Two fiber-optic cannulas (200-um core diameter) were implanted into bilateral dACC coordinates (±2 degrees angled, AP +1.2 mm, ML ±0.6 mm, DV −1.2 mm) for the stimulation of somata. Fiber optic cannulas (200-um core diameter) were implanted into ZI (bilateral; AP −1.8 mm, ML ±1.2 mm, DV −4.3 mm) or PnC (unilateral; AP −5.0 mm, ML +0.4 mm, DV −5.0 mm, not angled) coordinates for the experiments targeting projection terminals. For optogenetic control of PnC^GABA^ neurons, 300-nL solution of AAVdj-EF1a-DIO-hChR2-eYFP-WPRE-pA (1.2E+12 GC/mL) or AAV5-EF1a-DIO-eNpHR3.0-eYFP-WPRE-pA (6.1E+12 GC/mL GC/mL) or control AAVdj-syn-DIO-eGFP (2.3E+12 GC/mL) was injected into the PnC coordinate (as described above) in Vgat-ires-Cre transgenic mice. A fiber optic cannula with 200-um core diameter was implanted into the PnC (AP −5.0 mm, ML +0.4 mm, DV −5.0 mm, not angled).

For calcium activity monitoring of dACC→PnC neurons, 400-nL solution of AAV1-syn-FLEX-jGCaMP7s-WPRE (1.5E+13 GC/mL) was injected into the right (unilateral) dACC, and 200-nl solution of AAV retrograde-EF1a-Cre-WPRE-pA (GC/mL) was injected into the PnC, then a fiber-optic cannula (with 400-um core diameter) was implanted into the dACC coordinate (AP +1.2 mm, ML +0.5 mm, DV - 1.2 mm). For calcium activity monitoring of PnC^GABA^ neurons, 300-nL solution of AAV1-syn-FLEX-jGCaMP7s-WPRE (1.5E+13 GC/mL) was injected into the right PnC coordinate, and a fiber optic cannula with 400-um core diameter was implanted into the PnC.

For anterograde labelling of PnC neurons, 400-nL solution of AAV1-syn-Cre-WPRE-pA (3.4E+13 GC/mL GC/mL) was injected into the right (unilateral) dACC, and 300-nL solution of AAVdj-syn-DIO-eGFP (2.3E+12 GC/mL) was injected into the PnC. Eight weeks after injection, mice were sacrificed by transcardial perfusion for collection of the brain.

For retrograde tracing using CTB, 200-nL solution of a CTB tracer conjugated with Alexa 555 fluorescence was injected into the PnC. Ten days after injection, mice were sacrificed by transcardial perfusion for collection of the brain.

### Optogenetic stimulation

A 470-nm collimated diode and a 589-nm diode-pumped solid-state (DPSS) laser devices were used for the optogenetic activation and inhibition experiments, respectively. For photoactivation mediated by ChR2, 470-nm 20-Hz square pulses (5-ms pulse width, ~9 mW at the fiber tip) were delivered through the patch cord and the connected fiber-optic cannula. For photoinhibition mediated by eNpHR3.0, continuous yellow light (589-nm, ~6 mW at the fiber tip) was delivered through the fiber-optic components.

### Behavior with fiber photometry

#### Fiber photometry

A dual-wavelength fiber-photometry system (Doric Lenses) was assembled with a dichroic mini cube (iFMC4), 405-nm and 465-nm connectorized LEDs, a fluorescence detector/amplifier, then connected with a pyPhotometry controller board (1.0.2) and operated by Python script provided by pyPhotometry. Calcium-dependent fluorescence (465-nm excitation; F465) and isosbestic control fluorescence (405-nm excitation; F405) were monitored at 130-Hz sampling rate using 1-color time-division mode (alterations), and the data were analyzed by a custom-built LabView software. Isosbestic F405 fluorescence was fitted to F465 signals by least mean squares fitting (F405^fitted^). Motion-corrected fluorescence signal (dF/F) was calculated by: (F465 – F405^fitted^)/F405^fitted^. Both breathing monitoring (PowerLab) and fiber photometry systems received a synchronization signal (3.3-V or 5-V TTL) generated by a cDAQ output device (NI-9401, National Instruments) or a Raspberry Pi 4B device.

#### Drinking behavior

Prior to the experiment, mice were restricted from water for 24 hours. Mice were connected to an electric cord and an optical patch cord for the monitoring of breathing and calcium signals, respectively, and placed in a cylindrical chamber (15-cm height, 11-cm diameter). For one trial, a 10-ul water bolus (droplet) was provided through a small hole (~1.5 cm diameter) using a micropipette. Trials were repeated (with ~30 s inter-trial intervals) until the mouse refuses to drink. Behavior was video-recorded at a 25-Hz frame rate. The ‘drinking behavior’ for peri-event analysis (breathing and calcium signals) was defined by the moment (frame) the water bolus was absorbed into mouth.

#### Withdrawal and swimming

Before exposure to water (25°C water, 12-cm depth, placed in a glass cylinder with 15-cm diameter), electric and optical connections were thoroughly covered with a hydrophobic jelly (petroleum). Mice were lifted by their tail and slowly descended into the water until their lower face and upper limbs were submerged. For withdrawal experiment, mice were removed by lifting after 1-2 s of brief submersion. For release/swimming experiments, mice were gently released and allowed to swim for 30-60 s. Behavior was video-recorded from the side at a 10-Hz frame rate.

#### Shock-induced vocalization (squeaking)

Mice were connected to electric and optical patch cords (for breathing monitoring and photostimulation, respectively) and placed in a chamber equipped with a grid floor for shock delivery. Behavior was video-recorded from the top at a 10-Hz frame rate, and sound waveform was recorded at a 22 kHz sampling rate. During the experiment, mice received four repeats of electric foot shocks (2 s, 0.3 mA) with 30-60 s inter-trial intervals. The sound file (wav) was high pass filtered at 7 kHz to remove background noise, allowing for the extraction of the sound of squeaking vocalization.

#### Object sniffing

Mice were connected to electric and optical patch cords and placed in a shoebox cage with fresh cage beddings (29 × 18 × 13 cm). After an acclimation period of 3 min, a novel object (wooden cylinder) was placed in the cage at a location opposite to where the mouse was positioned. The mouse’s behavior was video-recorded from above at a 10-Hz frame rate. Three episodes of contact-sniffing behavior were analyzed for each mouse in conjunction with breathing cycles and calcium activities. The onset of sniffing was defined by the moment that the mouse contacted the object with its nostrils, whereas the termination of sniffing was defined by the moment of retraction behavior.

#### Elevated plus maze (EPM)

A custom-built plus-shaped maze made with white Plexiglass (77-cm long opposite closed arms, 77-cm long closed arms, and a square-shaped center plate with 7-cm sides) was situated 70 cm above the floor. For the EPM test, mice were connected to electric and optical patch cords, placed on the center plate, and allowed to voluntarily explore the environment for 10 min. Behavior was video-recorded from the top at a 10-Hz frame rate using a custom-built LabView software. For data synchronization, pyPhotometry and breathing monitoring (PowerLab) systems received a digital output (TTL) signal generated by cDAQ NI-9401 device (National Instruments; operated by the LabView software).

#### Elevated platform

A circular plastic platform (12-cm diameter) was equipped on a tripod and situated 100-cm above the ground. An opaque plastic tube (15-cm height, 11-cm diameter) was placed on top of the platform. Mice were connected to electric and optical patch cords and placed in the cylinder. After acclimation for 15 min, the tube was gently lifted by hand and mice were exposed to inescapable height. Behavior was video-recorded from the side at a 10-Hz frame rate.

### Behavior with optogenetics

#### Real-time place preference (RTPP)

A two-chamber white Plexiglass box with dimensions of 60 × 30 × 30 cm (W × D × H) was used for the RTPP test. Mice expressing ChR2-eYFP or eGFP constructs in dACC→PnC neurons were connected to a bilateral optical patch cord, placed in the testing arena, and allow to freely explore the environment for 30 min. Behavior was video-recorded from the top at a 10-Hz frame rate, and the location of the mouse (body center) was monitored in real time using EthoVision 11 tracking software (Noldus). During the second epoch (the second 10 min), one of the chambers was paired with photostimulation. The pairing of stimulated chamber (left or right) was counterbalanced for each mouse.

#### Female odor preference

Male mice (in the same groups used for RTPP) were used for a female odor preference test. Mice were connected to optical patch cord and placed in the same two-chamber testing arena. One side of the box (counterbalanced for each subject) was equipped with a 30-mm Petri dish that contains cage bedding collected from a female cage. Male mice were allowed to freely explore the environment for 30 min. During the second epoch (10 min), photostimulation was paired with the odor-equipped chamber.

#### TMT (Trimethyl-1,4,5-thiazoline) avoidance test

A plastic cage with dimensions of 47 × 26 × 21 cm (W × D × H) was used for the TMT avoidance test. Mice from the same groups (male mice, used for the RTPP and female odor test) were placed in the testing cage. One side of the cage (counterbalanced for each subject) was equipped with a 30-mm Petri dish that contains 300-uL of 10% TMT. Mice were allowed to freely explore the environment for 15 min. During the second epoch (5 min), photostimulation was delivered.

#### Elevated plus maze (EPM)

A plus-shaped white Plexiglass arena composed of two closed arms (77 cm), two open arms (77 cm), and a square center plate (7 cm) was situated 70 cm above the floor. For photoinhibition experiments with low baseline anxiety, mice were habituated by 5-10 min handling for 7 days. On the test day, mice were connected to optical patch cord then placed in the center plate of the EPM arena. Mice were allowed to explore the environment for 15 min. During the second epoch (5 min) of photoactivation experiments, 20-Hz 470-nm photostimulation was delivered. During the second epoch of photoinhibition experiments, continuous 589-nm light was delivered when mice were in center and open arms.

#### Looming-hide test

A plastic cage with dimensions of 47 × 26 × 21 cm (W × D × H) was used for the looming-hide test. Similar to a previous study^26^, the cage was equipped with a hiding shelter 13 × 24 × 21 cm (W × D × H) and an overhead display. Mice were acclimated in the testing chamber for 4-5 min and the time spent in the hiding shelter was monitored as baseline behavior. After acclimation, 15 repeats of expanding looming disk (black, 25-cm diameter) were presented in 24 s. Mice that successfully ran into hiding shelter received continuous light stimulation (20-Hz 470-nm) for 4 min.

#### Light/dark choice test

A white Plexiglass arena with dimensions of 41 × 41 × 30 cm (W × D × H) was equipped with an opaque cardboard shade (40 × 20 × 20 cm; W × D × H). The apparatus was placed under white fluorescence illumination and half the area (dark zone) was shadowed by the shade. Mice received surgical procedures to implant the nasal thermistor with the expression of AAV constructs for optogenetics. For photoinhibition experiments with low baseline anxiety, mice were habituated by 5-10 min handling for 7 days. A rotary joint (HRJ_OE_12_FC, Doric Lenses) with electric and optical connectors was used for resolving the entanglement between electric and optical patch cords. On the test day, mice were connected to the patch cords and placed in the testing arena, then allowed to freely explore the environment for 10 min. Photostimulation was delivered when mice were travelling the light zone throughout the 10-min testing.

#### Drinking test

Prior to the experiment, mice were restricted from water for 24 hours. Mice were connected to electric and optical patch cords (for breathing monitoring and photostimulation, respectively) and placed in a cylindrical chamber (15-cm height, 11-cm diameter). Before the testing session, mice were acclimated in the cylinder for 2 min with display of 10-ul water droplets. Then, in a 10-minute testing session, mice received 5 repeats of 20-s water displays without photostimulation (OFF session; baseline response), and then 12 repeats of the 20-s displays with photostimulation (ON session) in 5-30 s inter-trial intervals (ITIs). On the third session (OFF), mice were displayed with water droplets for 12 times without photostimulation. Behavior was video-recorded at a 25-Hz frame rate. Breathing cycles within the time frame of −100 to +500 ms from drinking were used for the analysis.

### Histology

#### Section preparation

Mice were transcardially perfused with phosphate buffered saline (PBS) then 4% paraformaldehyde dissolved in phosphate buffer (4% PFA/PB). Dissected brains were additionally fixated in 4% PFA/PB for 12-16 h and dehydrated in 30% sucrose/PB for 48 h, then sectioned coronally (40-um thickness) on a −20°C cryostat.

#### Immunohistochemstry

Brain sections covering the coordinate from AP −4.9 to −5.0 mm relative to the bregma were collected from three mice (3 sections from each, total 9 sections). Sections were incubated in blocking solution (0.3% Triton X-100 and 2% normal donkey serum, dissolved in PBS) for 1 h at room temperature, rinsed with PBST for three times, and incubated with rabbit anti-GAD67 (1:1000, dissolved in blocking solution; Merck Millipore SAB4300642) primary antibody for 16 h overnight. The following day, the sections were rinsed with PBST and incubated with Cy3-conjugated donkey anti-rabbit IgG (1:500, dissolved in blocking solution; Vector Laboratories) secondary antibody for 2 h. After rinsing with PBS, sections were mounted on slide glass and cover-slipped with DAPI Fluoromount-G solution.

#### Microscopy

Images were taken with a BZ-X710 all-in-one fluorescence microscope (Keyence). Region-wide images to confirm construct (AAV) expression were taken at 4× and 10× magnification. Precise images for observing axon terminals were taken at 20× magnification. Confocal microscopy images for co-localization analyses were taken using an FV3000 confocal laser scanning microscope (Olympus). Three to four images were taken from each brain section at 40× magnification. Serial Z-stack images were taken (6-12 planes, 1.0 aerial unit) in case cells were located on different focal planes.

### Statistical analysis

Statistical analyses were performed using GraphPad Prism 6. Normal distribution of dataset was tested by Shapiro-Wilk normality test (n ≥ 7) or Kolmogorov-Smirnov test (n < 7). In case the normality test failed (*P* < 0.05), non-parametric tests were performed. Repeated measures (RM) two-way ANOVA followed by Sidak’s post hoc test was used for between-group comparison with repeated measures. Two-tailed paired t test and Wilcoxon signed rank test were used for within-group analysis. Two-tailed unpaired t test and Mann-Whitney U test were used for between-group analysis. Details of statistical tests are described in the Extended Data Table 1.

**Extended Data Figure 1.**
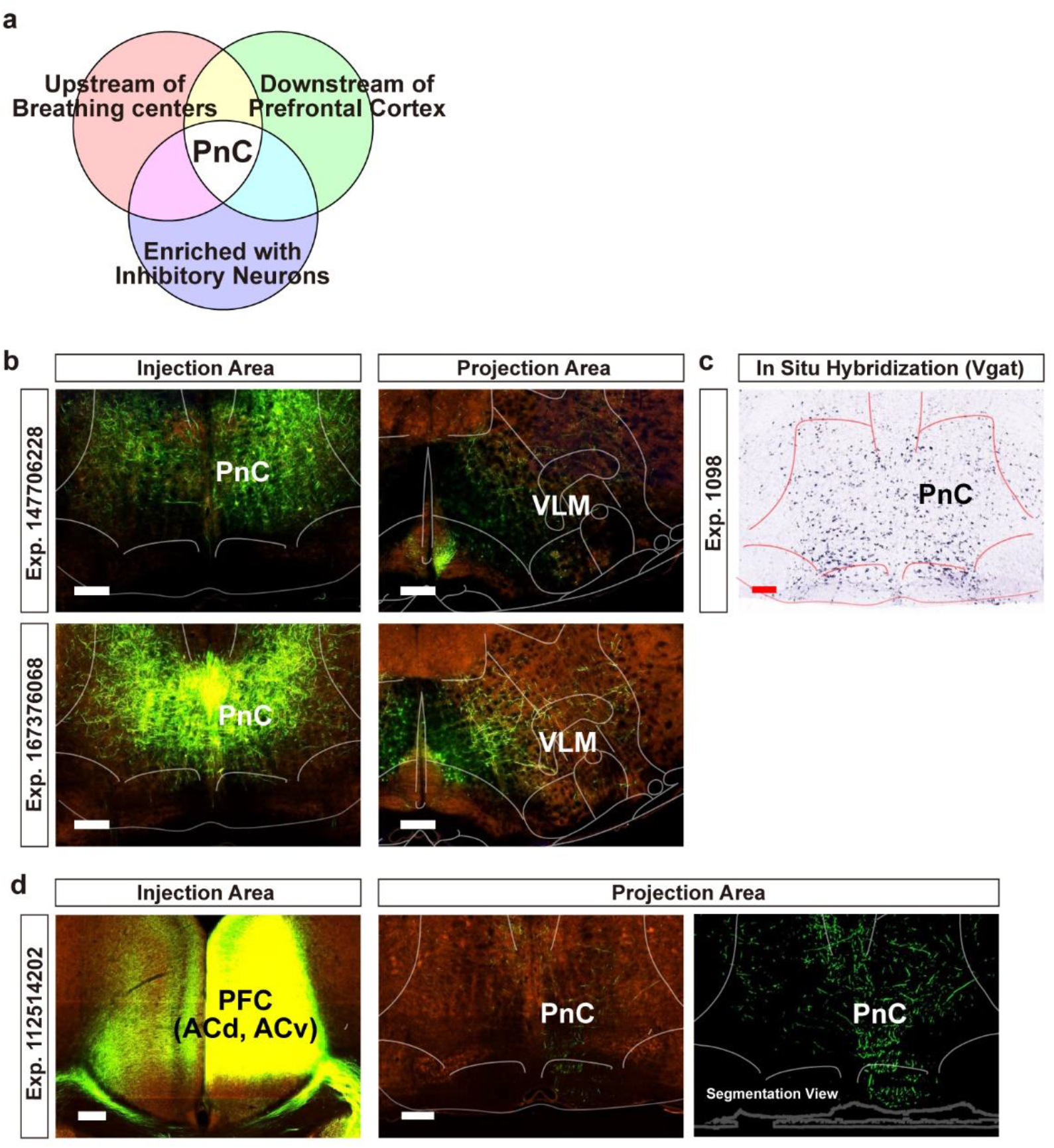
Searching for candidate circuits to mediate top-down inhibitory control of breathing based on the Allen Brain Atlas. **a**, Summary of the criteria used to search candidate circuits. Axonal projection images were searched through the Mouse Brain Connectivity Atlas (Allen Brain Institute). In situ hybridization (ISH) images were searched through the Mouse Brain Gene Expression Atlas (Allen Brain Institute). **b**, Sample images showing the projections of labelled PnC neurons observed in the ventrolateral medulla (VLM). Left images show the AAV injection site (PnC) with the expression of eGFP. Right images show the eGFP-labelled terminals in the VLM. **c**, Sample image from a Vgat ISH experiment. The PnC is enriched with neurons that express Vgat mRNA (GABAergic cell marker). **d**, Sample images showing the projections of PFC neurons observed in the PnC. Left image shows the AAV injection site (PFC, prefrontal cortex; covering the dACC and vACC) with the expression of eGFP. Middle and right (in segmentation view) images show the eGFP-labelled axon terminals in the PnC. Scale bar, 200 μm.

**Extended Data Figure 2.**
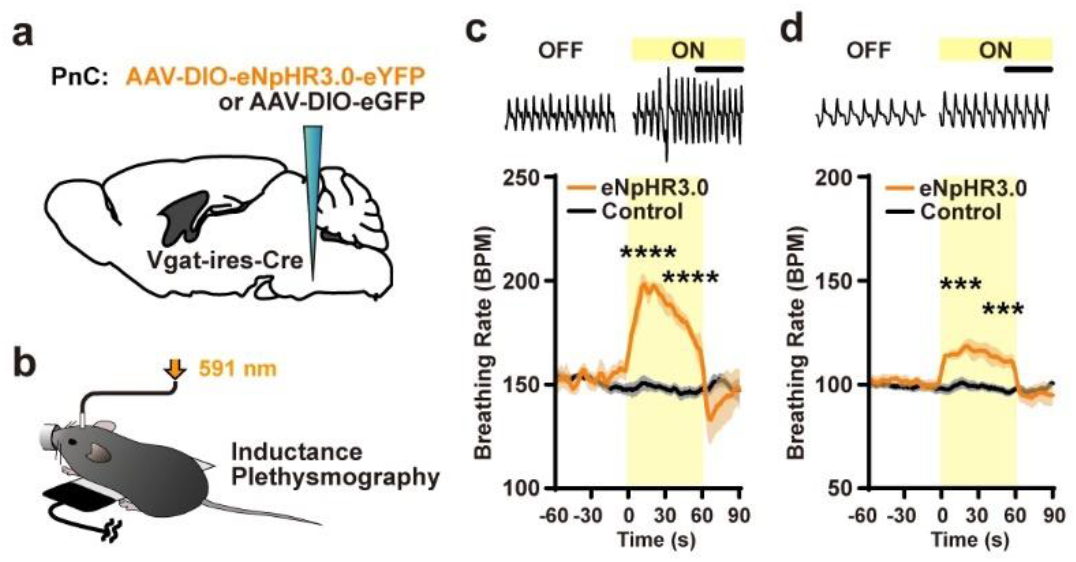
Photoinhibition of PnC^GABA^ neurons increases breathing rate. **a**, Schematic of the virus injection surgery. **b**, Breathing signals were monitored by inductance plethysmography with the delivery of 591-nm light for photoinhibition. **c**-**d**, Photoinhibition of PnC^GABA^ neurons induced increased breathing rate in the eNpHR3.0 group. Top, raw breathing signals. Bottom, average (mean) breathing rate in eNpHR3.0 (n = 4) or Control (eGFP, n = 3) groups. Two different baselines at 150 BPM or 100 BPM were used in **c** and **d**. Statistical analysis was performed with repeated measures (RM) two-way ANOVA followed by Sidak’s post hoc test. Between-groups comparison ***p < 0.001, ****p < 0.0001. Data are shown as mean ± s.e.m.

**Extended Data Figure 3.**
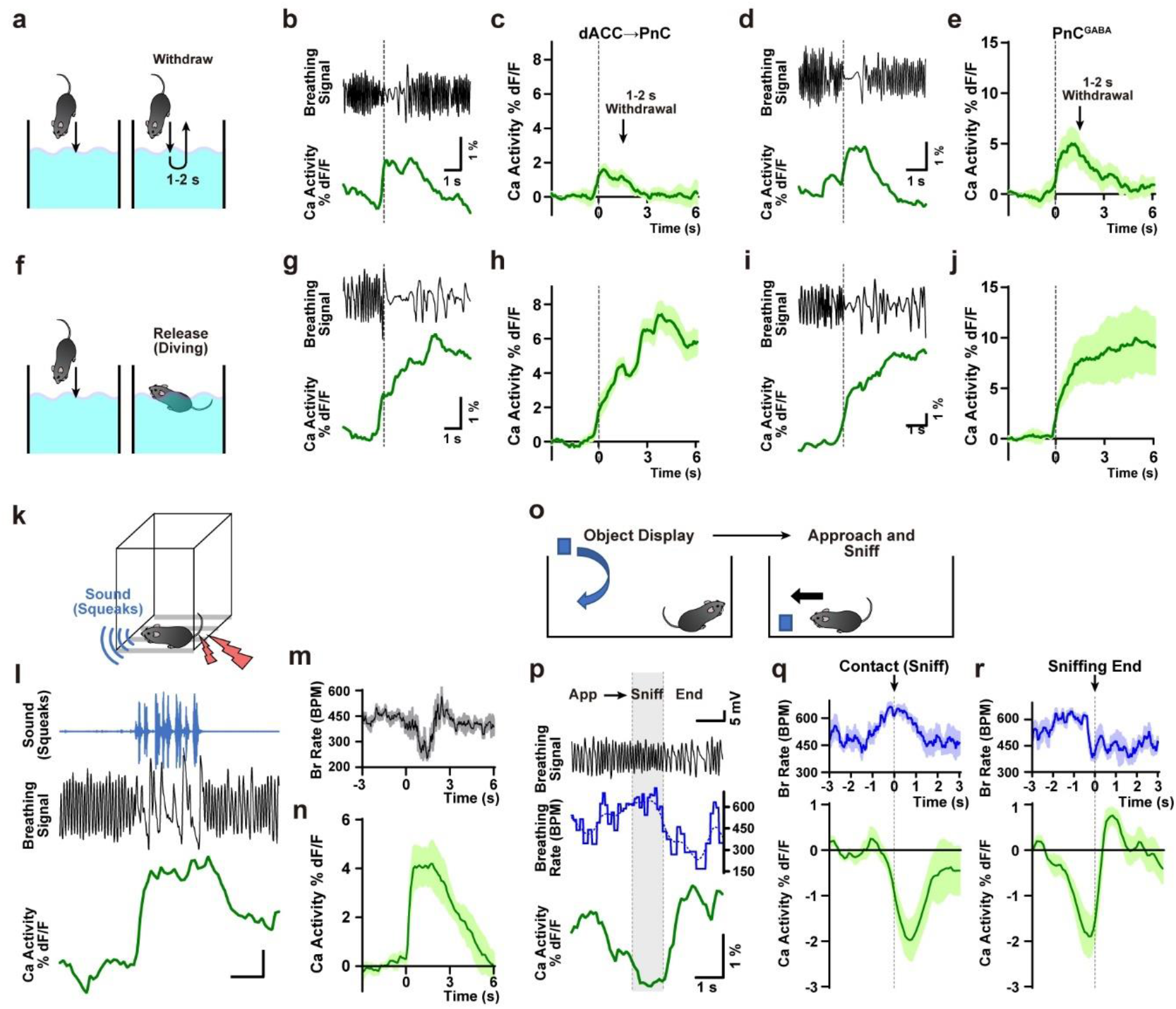
dACC→PnC and PnC^GABA^ calcium activity during swimming, squeaking, and sniffing behaviors. **a**, Schematic of “withdrawal” experiment in which mice were briefly (1-2 s) submerged in water and then removed. **b-e**, Monitoring results from mice expressing jGCaMP7s in dACC→PnC neurons (**b**, representative trace; **c**, averaged n = 6) or PnC^GABA^ neurons (**d**, representative trace; **e**, n = 5) with representative breathing traces (**b**, **d**). **f**, Schematic of “release” experiment in which mice were released after submersion. **g-j**, Monitoring results from mice expressing jGCaMP7s in dACC→PnC neurons (**g**, representative trace; **h**, n = 5) or PnC^GABA^ neurons (**i**, representative trace; **j**, n = 5) during release/swim experiment. **k**, Schematic of pain-induced vocalization (squeaking experiment). **l**, Representative traces showing sound waveform (squeaks; top), breathing cycles (middle) and dACC→PnC calcium activity (bottom). **m**, Averaged breathing rate was decreased during the pain-induced vocalization. **n**, Averaged calcium activity trace from dACC→PnC neurons during pain-induced vocalization (n = 4). **o**, Schematic of object sniffing experiment. **p**, Representative traces showing raw breathing cycles (top), breathing rate (middle), dACC→PnC calcium activity. **q**, At onset of contact-sniffing events, fast breathing rate (~600 BPM; top) and subsequent decrease of calcium activity was observed (bottom; n = 4). **r**, At the end of sniffing events, decreased breathing rate (top) and rebound-like increase of calcium activity was observed (bottom; n = 4). Data are shown as mean ± s.e.m.

**Extended Data Figure 4.**
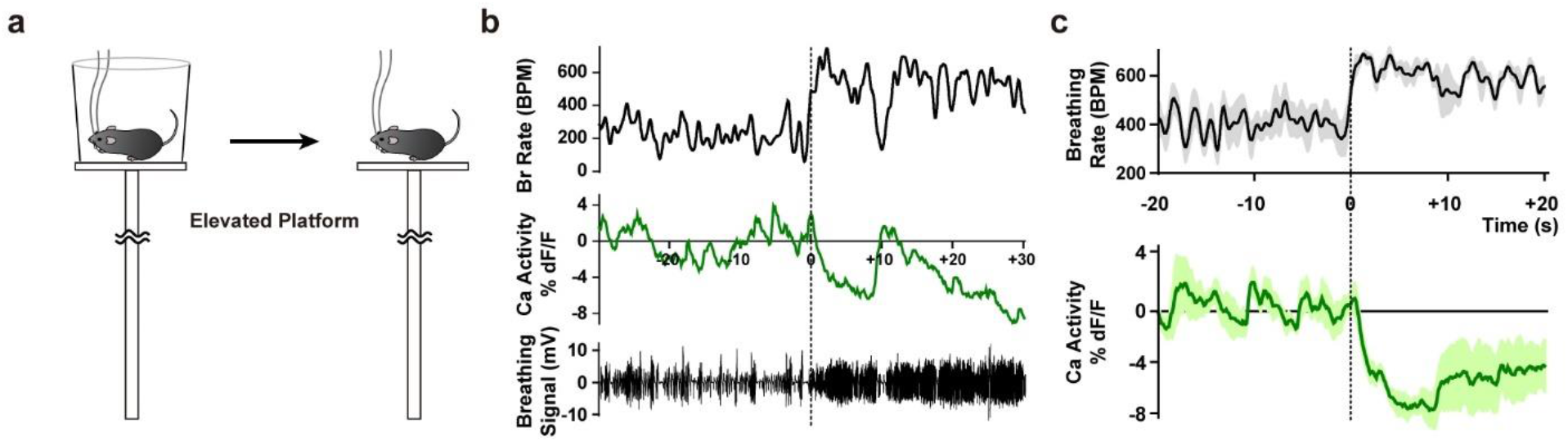
Breathing and dACC→PnC activity monitored on an elevated platform. **a**, Schematic of the elevated platform experiment. Mice were initially placed in an opaque plastic cylinder, then exposed to inescapable height by removing the cylinder. **b**, Representative traces showing breathing rate (top), raw cycles (bottom) and dACC→PnC activity (middle) changes after height exposure. **c**, Averaged traces (n = 5 mice) showing the breathing rate and dACC→PnC activity changes after height exposure.

**Extended Data Figure 5.**
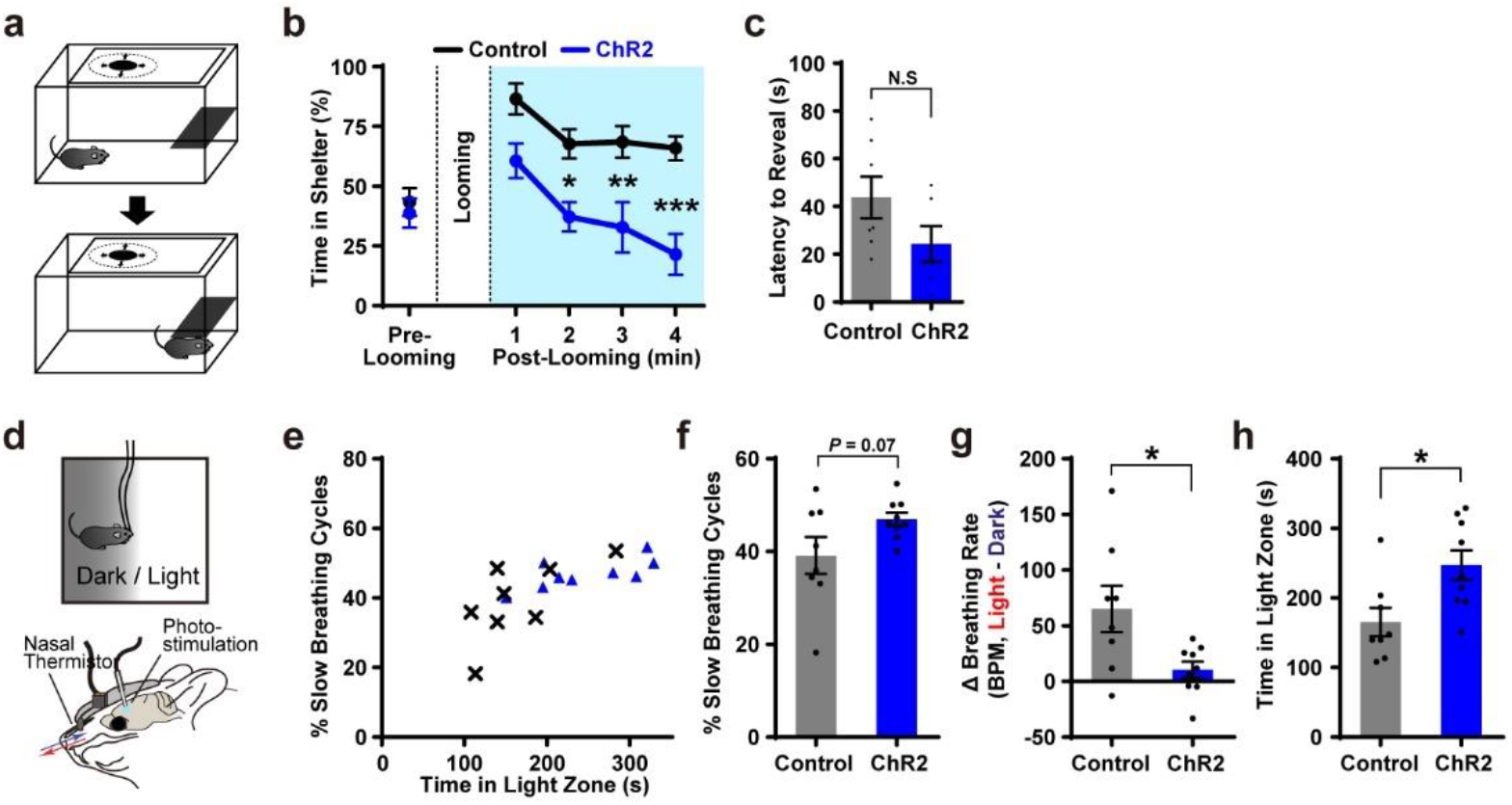
Photoactivation of dACC→PnC neurons alleviates anxiety-like behaviors. **a**, Schematic of looming-hiding experiment. Time spent in hiding (in shelter) was observed after overhead looming stimulus. **b**, Photoactivation of dACC→PnC neurons reduced time spent in hiding in the ChR2 group (n = 6), compared to the control group (n = 7). **c**, Latency to the first reveal was not affected by photoactivation. **d**, Schematic of light/dark choice test with breathing monitoring and photostimulation. **e-h**, Breathing and behavioral correlates observed in the light/dark choice test. **f**, Percent of slow breathing cycles in light zone (slower than median breathing rate). **g**, Difference of breathing rates between light and dark zones (ΔBPM, light minus dark). **h**, Time spent in light zone. **b**, Repeated measures (RM) two-way ANOVA followed by Sidak’s post hoc test. **c**, **f**, **g**, Mann-Whitney U test. **h**, Unpaired t test. Between-groups comparison *p < 0.05, *p<0.01 ***p < 0.001. Data are shown as mean ± s.e.m.

**Extended Data Figure 6.**
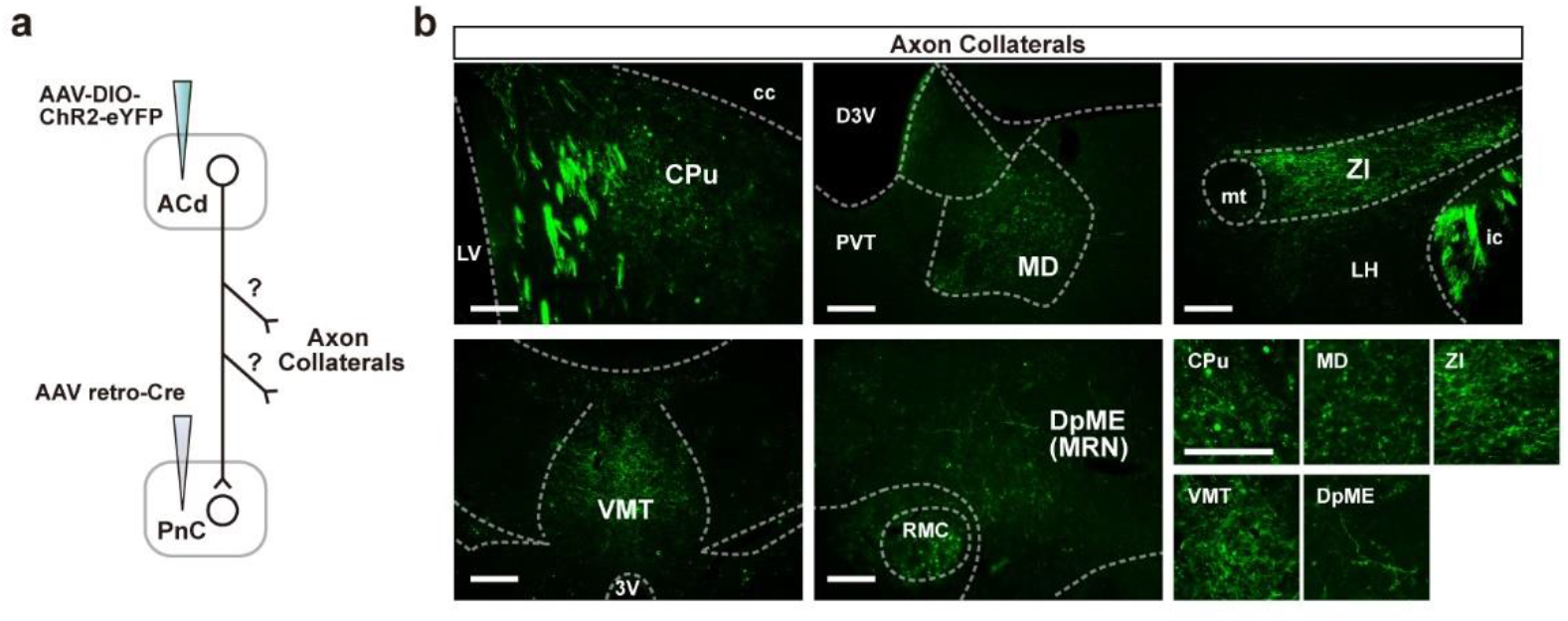
Axon collaterals of dACC→PnC neurons. **a**, Observation of axon collaterals. AAVretro-Cre was injected into the PnC and AAV-DIO-ChR2-eYFP was injected into the dACC. Putative axon collaterals were revealed by observation of eYFP-labelled axons in brain regions other than the PnC (AAVretro-Cre injection site) area. **b**, Images showing axon collaterals of dACC→PnC neurons. CPu, caudate putamen. cc, corpus callosum. LV, lateral ventricle. D3V, dorsal third ventricle. PVT, paraventricular nucleus of thalamus. MD, mediodorsal nucleus of thalamus. ZI, zona incerta. mt, mammillothalamic tract. LH, lateral hypothalamus. ic, internal capsule. VMT, ventromedial nucleus of thalamus. RMC, red nucleus, magnocellular part. DpME, deep nucleus of the midbrain. Scale bar, 200 μm.

**Extended Data Figure 7.**
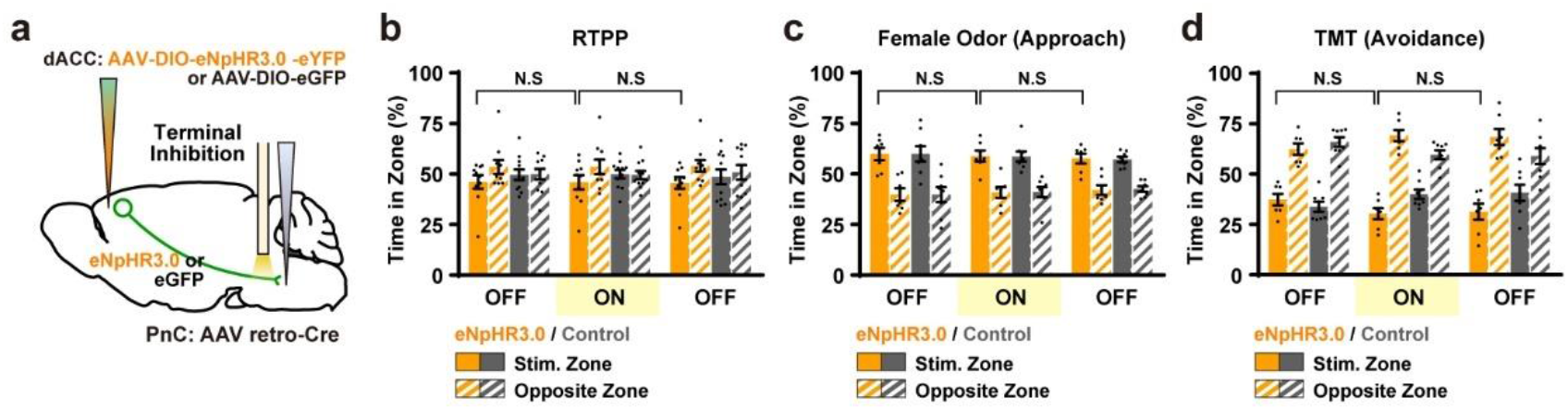
Real-time place preference, female odor preference, and TMT odor avoidance tests with Photoinhibition of dACC→PnC terminals. **a,** Schematic showing photoinhibition of dACC→PnC terminals. **b-d**, Behaviors in the RTPP test (**b**; eNpHR3.0, n = 10; eGFP, n = 11), approach response to female odor (**c**; eNpHR3.0, n = 7; eGFP, n = 8), and avoidance response to TMT (**d**; eNpHR3.0, n = 7; eGFP, n = 8) were not affected by the photoinhibition.

**Extended Data Figure 8.**
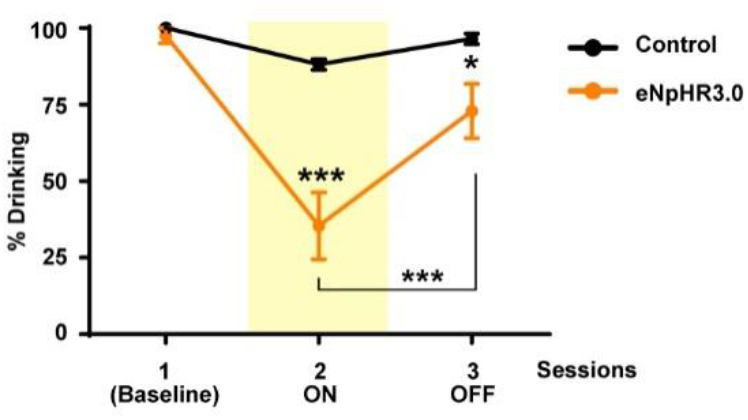
Percent of successful drinking responses throughout testing sessions with or without photoinhibition of dACC→PnC terminals. Repeated measures (RM) two-way ANOVA followed by Sidak’s post hoc test. Data are shown as mean ± s.e.m.

**Extended Data Table 1.**
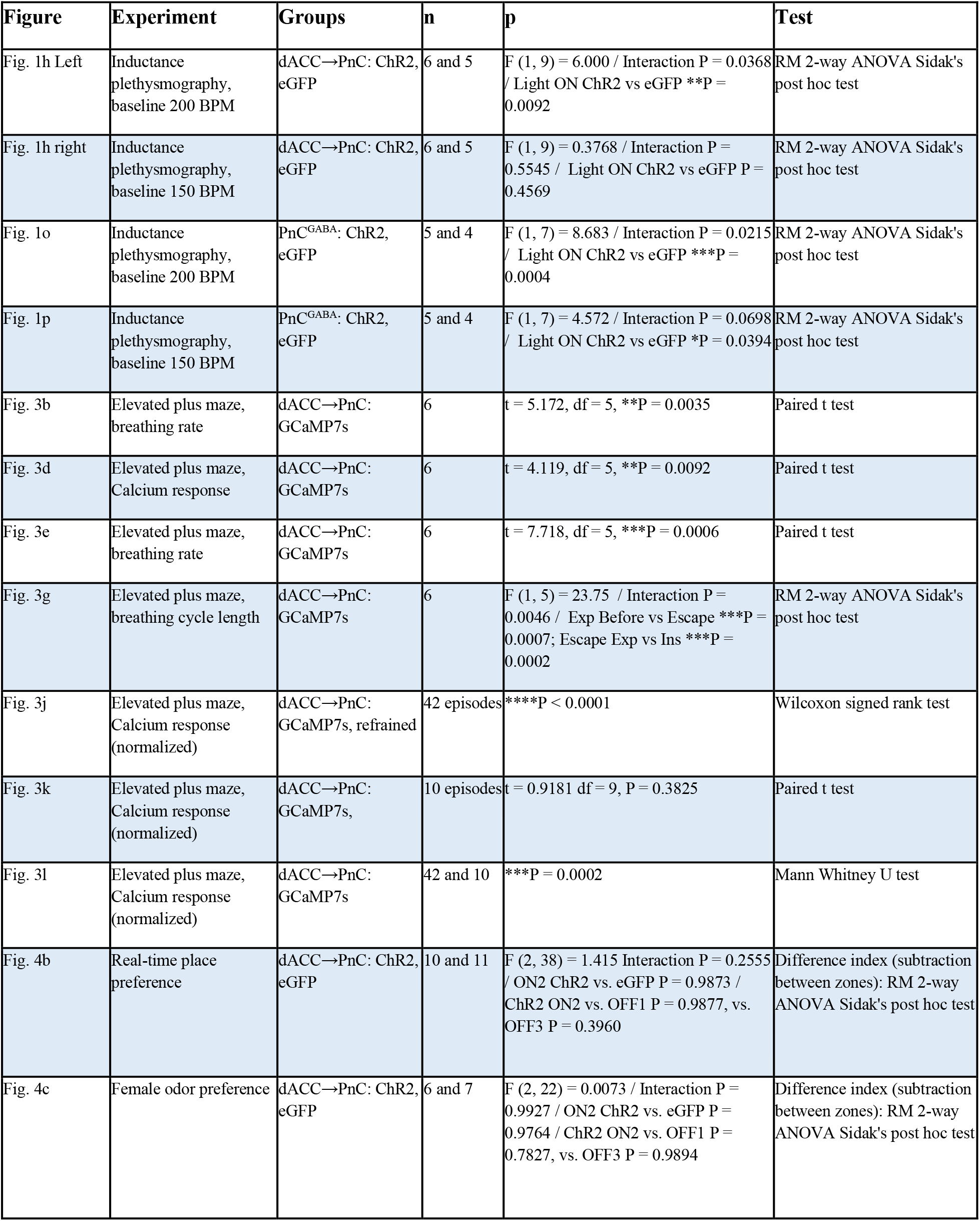
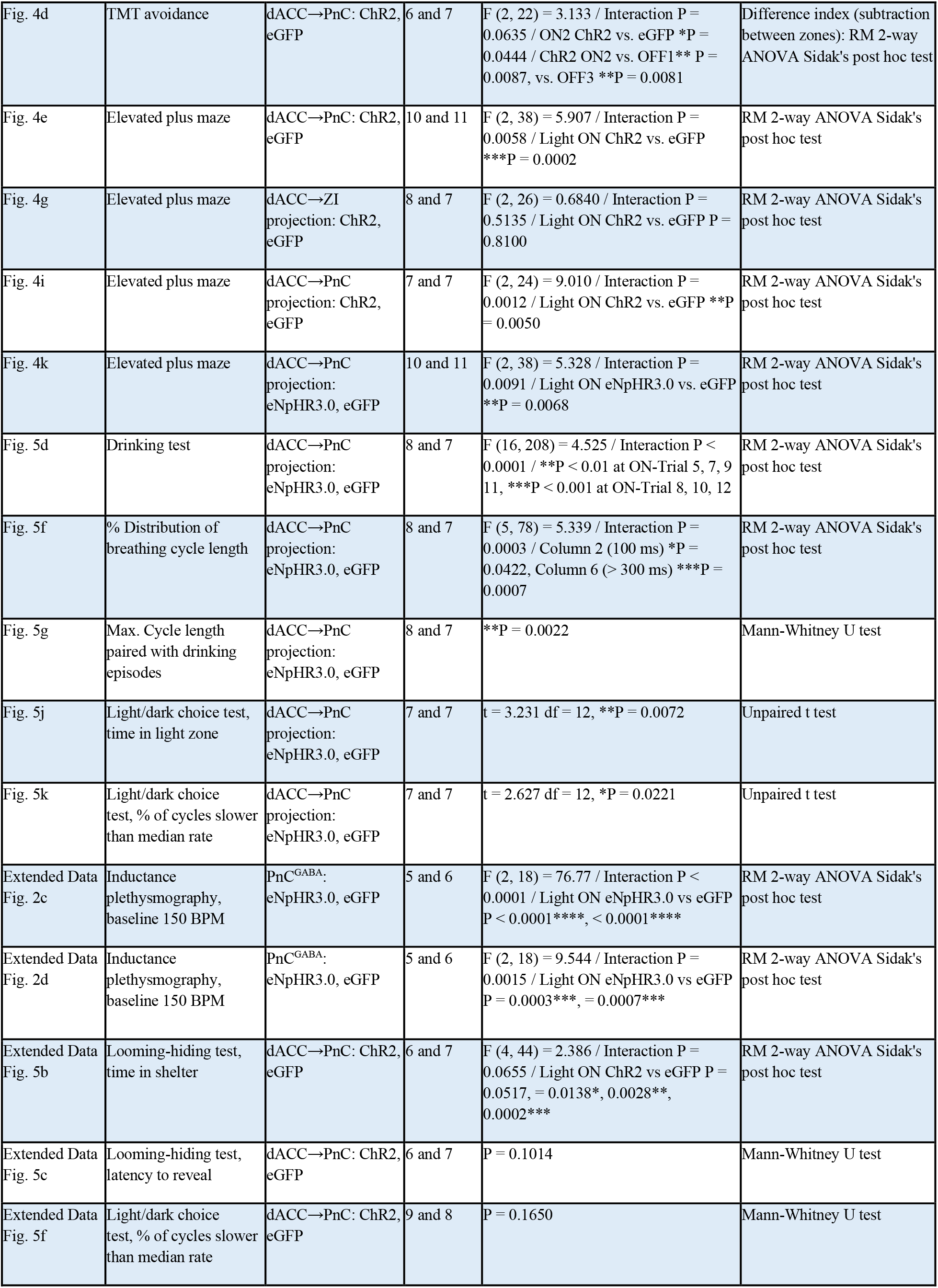
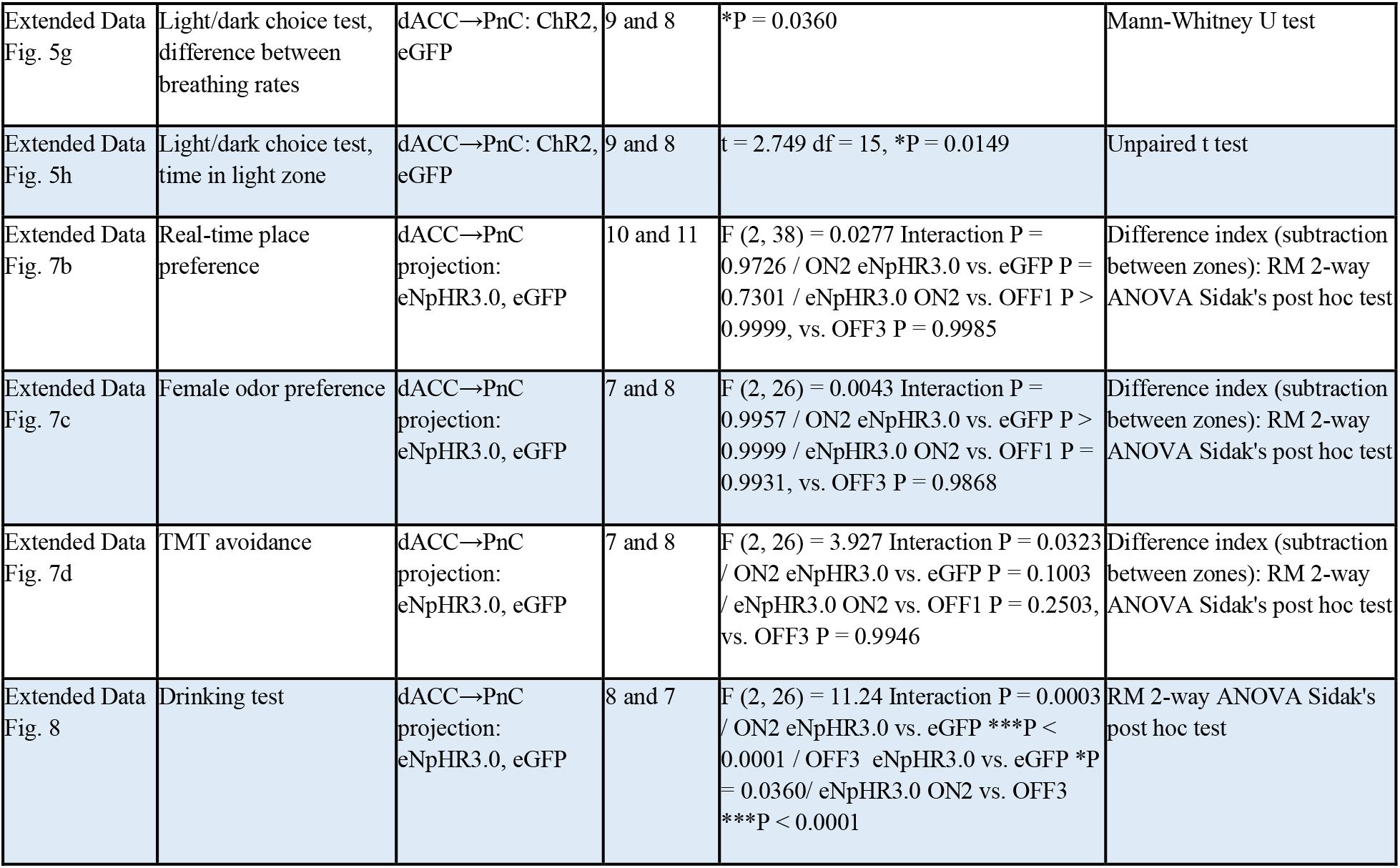
Statistical analyses

